# Intravital Multimodal Imaging of Human Cortical Organoids for Chronic Stroke Treatment in Mice

**DOI:** 10.1101/2025.07.05.663297

**Authors:** Jinghui Wang, Guanda Qiao, Honglin Tan, Mengyang Jacky Li, Kexin Wang, Jiadi Xu, Miroslaw Janowski, Tian-Ming Fu, Piotr Walczak, Yajie Liang

## Abstract

Chronic stroke leads to enduring neurological deficits and remains a major clinical challenge. Human induced pluripotent stem cell (hiPSC)-derived cortical organoids (COs) offer promise for regenerative therapies, yet their application in chronic stroke remains understudied, in part due to limited tools for monitoring graft in vivo. Here, we present a Multimodal Imaging Platform for Organoid Tracking (MIPOT), which integrates MRI, bioluminescence imaging, light microscopy, and two-photon fluorescence microscopy to noninvasively track transplanted COs in the post-stroke brain over time. Using MIPOT, we verified precise delivery of COs into cleaned stroke cavities, observed progressive declines in viability within the first 10 days, and introduced labeling methods for functional in vivo tracking of COs at subcellular resolution. By 4 weeks post-transplantation, histological analysis revealed survival of COs in the cerebral cortex. Notably, COs engrafted into the hippocampus displayed enhanced maturation, underscoring the role of local microenvironments in graft integration. MIPOT enables dynamic, noninvasive evaluation of hiPSC COs in chronic stroke, providing a foundation for mechanistic studies and translational development of organoid-based therapies.

## Introduction

Stroke is a leading cause of serious long-term disability^1^. Despite notable advancements in acute stroke management, thrombolytic therapy and mechanical thrombectomy are still time-sensitive (4.5-6 hours), and applicable to a limited number of stroke patients^2,3,3^. As most return-of-function after stroke occurs early and recovery plateaus 3 to 6 months after stroke onset^4^, chronic stroke, defined as 6 months to years after onset^5^, has a very high chance of leading to permanent disability ^6,7^. Therefore, there is an unmet need for more effective therapeutic strategies that can address the complex, established neurological deficits of chronic stroke. Unfortunately, current therapeutic approaches have not yielded significant benefits for patients with chronic stroke, despite the immense efforts spent^8,9^.

Human induced pluripotent stem cell (hiPSC)-derived brain organoids are self-organizing, three-dimensional (3D) structures that recapitulate key aspects of brain development, including the formation of region-specific architectures such as the midbrain, hippocampus, and cerebellum^10–12^. Notably, the brain organoids show 3-dimensional neural connectivity and brain functionality that recapitulate features of brain development and maturation ^12,13^. Compared to the transplantation of isolated neural stem or progenitor cells, brain organoid transplantation offers several advantages for neural repair. One major advantage is that they contain a structured microenvironment composed of diverse neural cell types—including progenitors, neurons, astrocytes, and oligodendrocytes^14,15^—that support each other and enhance graft survival following transplantation ^16^. In addition, whereas transplantation of a single neural stem or progenitor cell type may be insufficient to regenerate all components of damaged brain tissue ^17,18^, organoids provide a rich and varied cellular source capable of contributing to more comprehensive tissue reconstruction. Furthermore, brain organoids can be customized to mimic specific brain regions, allowing for the development of targeted therapies based on the anatomical and functional needs of the damaged area ^16,19,20^. Together, these properties make brain organoids transplantation a promising approach for repairing or replacing damaged brain tissue for traumatic brain injury ^21^ and a range of neurodegenerative diseases ^16,19,22^.

Recent studies have explored the transplantation of hiPSC-derived cortical organoids (COs) for treating ischemic stroke in rodent models, demonstrating promising therapeutic effects ^23,24^. However, two major challenges remain. First, the dynamic behavior of transplanted organoids is poorly understood due to the absence of intravital imaging tools. Most previous studies have relied on static histological analyses at terminal time points, providing only limited snapshots of graft positioning, survival, and cellular behavior. Second, it remains unclear whether brain organoid transplantation can be effectively applied to chronic stroke, as prior studies have primarily focused on the acute or subacute phases. To address these gaps, we developed a robust intravital multimodal imaging platform and applied it to study CO transplantation during the chronic phase of stroke in a mouse model. By integrating Magnetic Resonance Imaging (MRI), bioluminescence Imaging (BLI), bright field microscopy and Two-Photon Fluorescence Microscopy (TPFM), our multimodal imaging platform for organoid tracking (MIPOT) enables longitudinal, multi-scale monitoring of graft location, viability, and morphological changes. We generated human COs via directed differentiation of hiPSCs and transplanted them into the infarct cavity following the surgical removal of the ischemic core at chronic phase of stroke. Remarkably, the engrafted COs survived and exhibited obvious differentiation into neurons when they are transplanted in host hippocampus. Together, this study establishes a novel framework for evaluating brain organoid therapies in chronic stroke and underscores the value of multimodal imaging to track and quantify post-transplantation graft status, ultimately advancing the clinical translation of organoid-based treatment for chronic stroke.

## Results

### Multimodal imaging platform for peri- and post-transplantation tracking of COs

To enhance the consistency, reliability, and quantitative assessment of organoid transplantation, we propose the Multimodal Imaging Platform for Organoid Tracking (MIPOT). MIPOT integrates four complementary intravital imaging modalities (surgical microscopy, MRI, BLI and TPFM) to monitor human iPSC-derived COs transplanted into the chronically ischemic brain (**Fig. 1**). Following photothrombotic stroke induction using Rose Bengal (RB), cortical infarcts are allowed to mature over a 10-day period to model chronic stroke before CO transplantation. MIPOT enables imaging across key phases of the transplantation workflow. At the time of transplantation, white-light microsurgical microscopy permits visualization of the ischemic core, its removal and graft delivery. In the post-transplantation phase, MRI confirms graft presence, location, and cavity filling and integration; BLI quantifies graft viability over time; and TPFM resolves subcellular morphology, migration, and functional integration. At the terminal endpoint, histological and immunohistochemical analyses validate graft fate and interactions with the host. Together, MIPOT enables systematic and dynamic evaluation of organoid engraftment and integration in a chronic stroke setting.

**Fig. 1:**
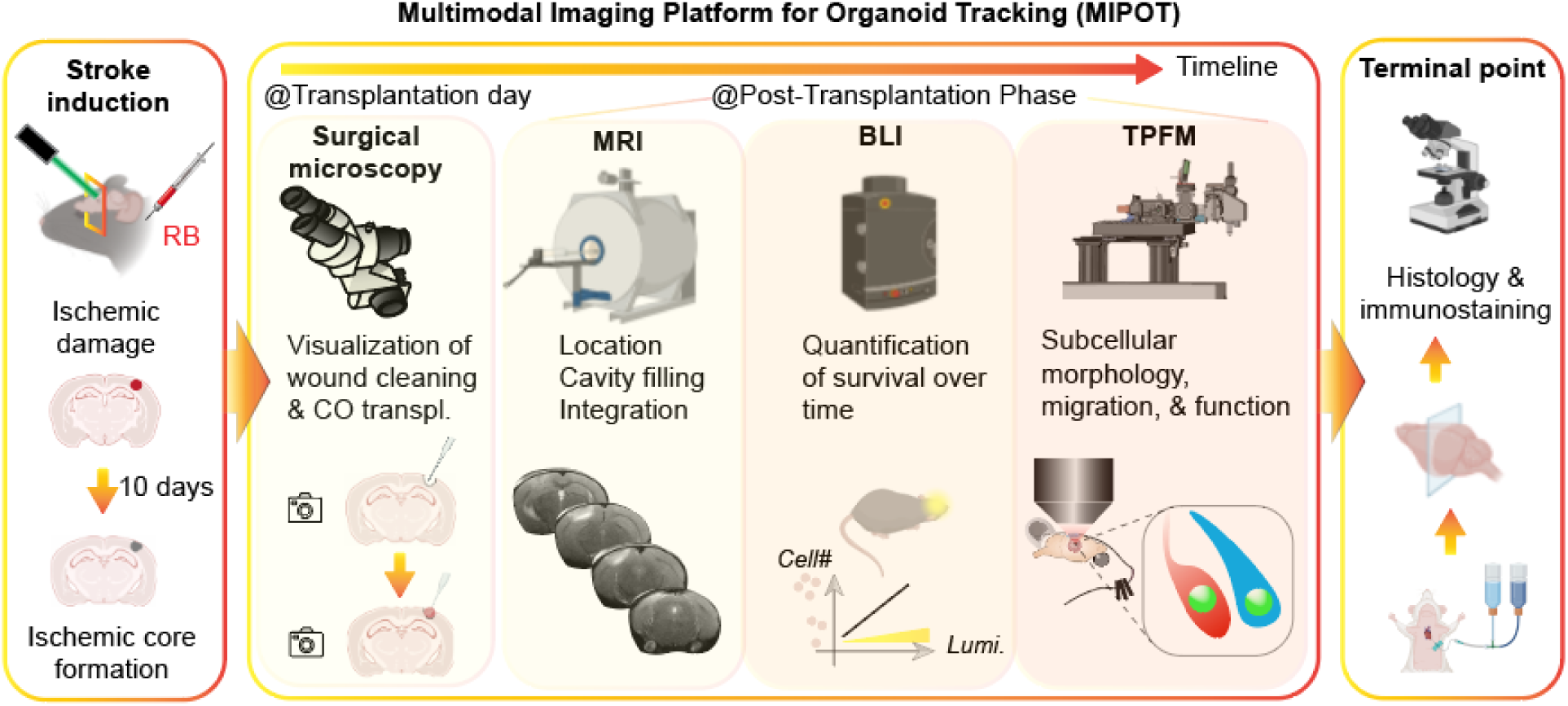
Schematic overview of MIPOT, a longitudinal imaging platform to monitor human CO transplantation in a mouse model of chronic stroke. Stroke is induced via targeted photothrombosis, and transplantation is performed 10 days post-stroke. Surgical microscopy guides intraoperative delivery. MRI confirms graft location and cavity filling. BLI tracks graft viability and survival, while TPFM enables high-resolution assessment of organoid morphology, migration, and integration. Terminal histology provides cellular and molecular validation.

### Generation of iPSC-COs and their transplantation for the treatment of chronic stroke

To generate COs enriched in cortical neuron subtypes, we adapted and extended a stepwise differentiation protocol using small molecules and neurotrophic factors (**Fig. 2a**). Human iPSCs were first characterized by immunostaining of the featured TFs (**Supplementary Fig. 1a**), then they were exposed to dual-SMAD inhibitors (Dorsomorphin and SB-431542) to induce neuroectodermal fate, followed by FGF2 and EGF to promote expansion of neural progenitors. From day 25, cortical specification and maturation were supported by BDNF and NT-3. Organoids maintained in this culture condition over 8 weeks exhibited characteristic changes in morphology, including size expansion and increased tissue density (**Fig. 2a**, lower panels and **Supplementary Fig. 1b-c**). At day 56 (D56), immunostaining revealed robust neuronal differentiation as evidenced by widespread expression of βIII-tubulin (Tuj1) across the organoid (**Fig. 2b**). Higher magnification views showed processes of Tuj1+ cells (**Fig. 2b**, arrows from insets), suggesting early cortical-like architecture. By day 80 (D80), organoids developed multilayered structures enriched with deep-layer cortical markers CTIP2 and upper-layer marker SATB2 (**Fig. 2c**), indicating progressive maturation and laminar organization.

**Fig. 2:**
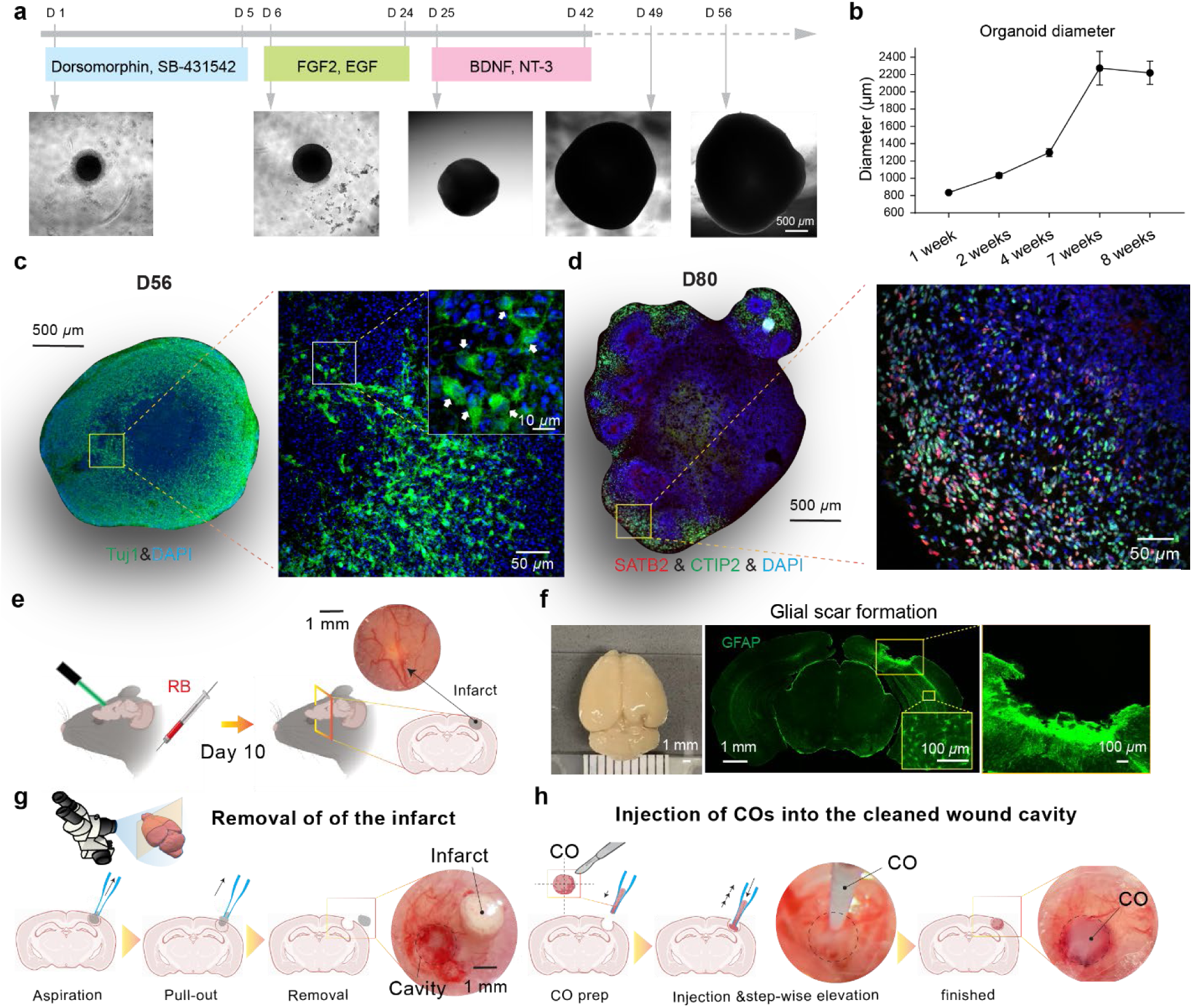
Generation of COs and the transplantation procedure for treatment of chronic stroke. **a** Timeline for CO differentiation. HiPSCs were differentiated into COs over 8 weeks using a step-wise factor treatment. Representative brightfield images show organoid morphology at different time points. **b** Change of organoids diameter over time (n = 6 - 12 COs per timepoint, Mean and STD was plotted)**. c** Immunohistochemistry of mature COs at D56 stained for Tuj1 (green, neuronal marker) and DAPI (blue, nuclei). The magnified inset (right) shows neuronal morphology and density, with white arrows indicating neuronal processes. **d** Spatial organization of cortical layers within D80 CO. Slices of D80 organoid (left) stained for SATB2 (purple, cortical layer II/IV marker), CTIP2 (green, cortical layer V/VI marker), and DAPI (blue, nuclei). The magnified inset (right) illustrates the presence and spatial arrangement of distinct cortical neuron populations. **e** Schematic illustrating the induction of an ischemic infarct by injecting Rose Bengal (RB) and illuminating the brain with a 532 nm laser. The inset shows the visible infarct region in the mouse brain. Representative image of a mouse brain (left) and a brain slice showing the robust glial scar formation around the infarct site immunostained with GFAP antibody 10 days post onset of stroke. **g** Schematic depicting the aspiration procedure to create a clean wound cavity after infarct formation under direct visualization through the microsurgical microscope. The inset shows the resulting cavity and dissected infarct tissue from the mouse brain under a surgical microscope. **h** Schematic illustrating the surgical procedure for injecting COs into the prepared wound cavity in the mouse brain. The subsequent images show the organoid successfully placed within the cavity.

To model chronic stroke, we employed a photothrombotic cortical infarction paradigm using RB^25,26^, followed by a 10-day infarct maturation period to establish infarcted tissue with glial scar formation which were visually confirmed under surgical microscope by a pale regions close to the cortical surface (**Fig. 2d**). Immunohistochemistry at day 10 post-stroke showed robust GFAP+ glial scar formation surrounding the lesion cavity (**Fig. 2e**). This defined injury stage provided a permissive window for surgical intervention. We then established a two-step transplantation procedure in the chronic stroke model under the direct visualization of surgical microscope. First, necrotic infarct tissue was removed via stereotaxic aspiration, exposing a well-defined wound cavity (**Fig. 2f**). Subsequently, COs at the age of D56 post-differentiation were first split into four pieces (each has a diameter of ∼ 1 mm) and then gently aspirated into glass pipettes (∼250 *μ*m inner diameter opening). Then they were slowly injected into the cleaned infarct space while step-wise elevation was performed as we previously described^27^ (**Fig. 2g**). This surgical approach enabled controlled and reproducible delivery of COs into the lesion site, forming the foundation for assessing post-transplantation integration and repair.

### In vivo MRI monitoring of transplanted COs

To assess initial engraftment and monitor the spatial characteristics of transplanted COs in vivo, we employed high-resolution T2-weighted (T2w) MRI following transplantation using a 9.4T MRI scanner (Bruker, Ettlingen, Germany). MRI scans were acquired at −3, 1 and 14 days post-transplantation of COs to capture dynamic changes in graft location and property (**Fig. 3a**). T2 hyperintensities corresponding to the injection site allowed clear visualization of cavity filling (**Fig. 3b**) and provided a noninvasive means to confirm successful graft delivery. Organoids appeared as distinct T2 hyperintensity regions within the infarct cavity on Day 1 post-transplantation, confirmed by over-head photos from light microscopy (dashed regions, **Fig. 3b**). On Day 14, the border of COs was not as obvious as on Day 1. To find out the exact border of the graft, we performed immunostaining of CO through human antigen marker (STEM121) and registered the same coronal brain slice (slice#4, **Fig. 3b**) under MRI and histology (**Fig. 3c**), which clearly reveals the outline of implanted CO as well as the viable grafted cells (**Fig. 3f**, inset). Interestingly, the pattern of inhomogeneous T2 hypointensity regions in the CO engraftment site (**Fig. 3d**) matched well with the histology observation (**Fig. 3e-f**): dark spots colocalize with densely packed nuclei with fewer human cells (arrows in **Fig. 3d** and **3f**).

**Fig. 3:**
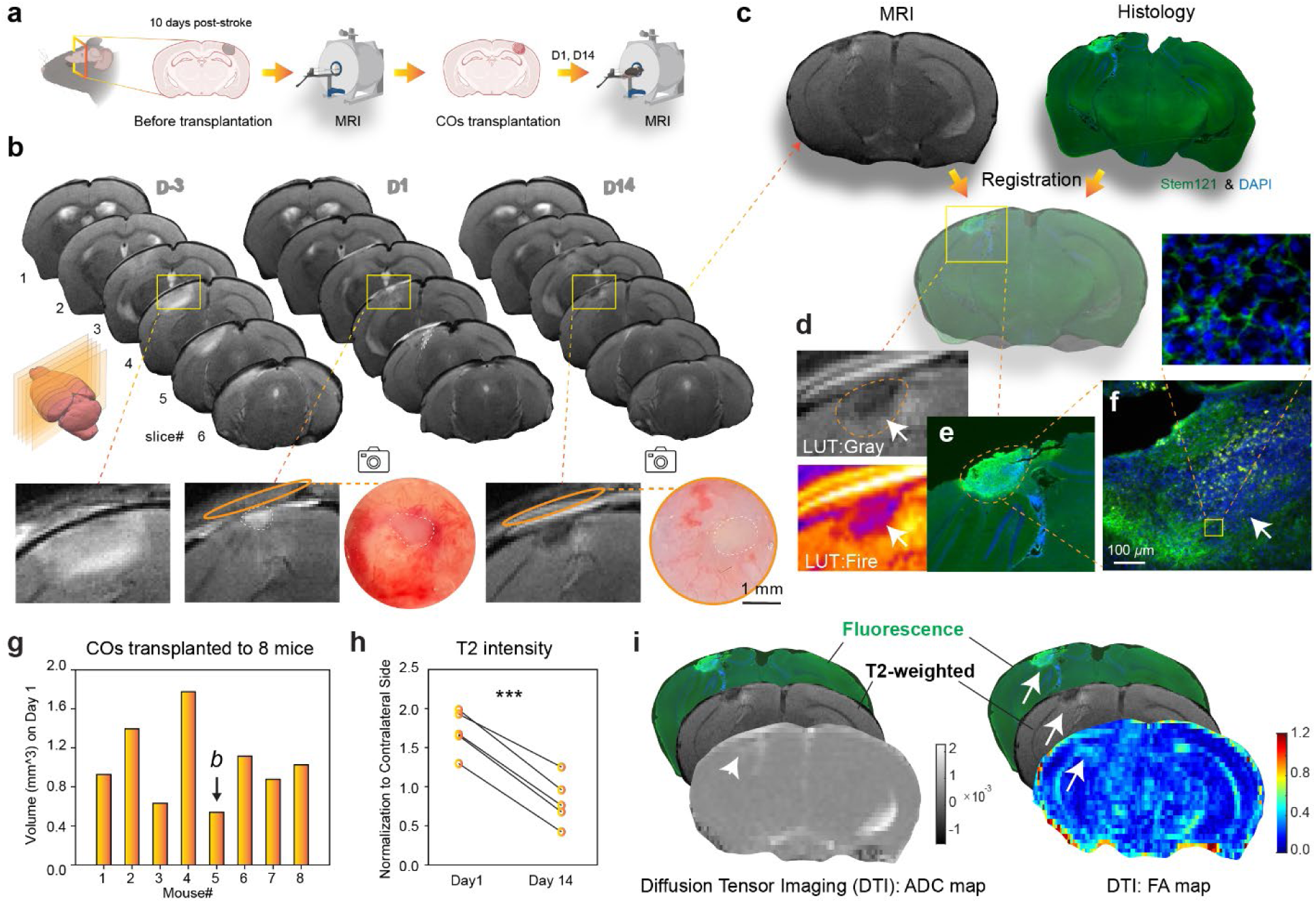
In vivo MRI monitoring of transplanted COs. **a** Schematic illustration of the experimental timeline for MRI scans and CO transplantation. **b** Representative T2-weighted (T2w) MRI images of a mouse brain at D-3, D1, and D14. Yellow boxes indicate the magnified regions shown in the top rows. The bottom row displays overhead light microscopy photos of the corresponding brain regions, confirming the presence of the transplanted COs. **c** Co-registration of MRI and histology images. The T2w MRI image (left) and the corresponding histology image stained for human antigen marker (STEM121, green) and DAPI (blue) (right) are shown for slice #4 at D14. **d** Magnified view of the CO engraftment site from the T2w MRI on D14, showing inhomogeneous T2 hypointensity regions (top, LUT Gray) and corresponding pseudocolored image (bottom, LUT Fire). **e** Corresponding histological image of the CO engraftment site, showing the outline of the implanted CO and viable grafted cells. **f** Higher magnification of (**e**) with arrows indicating areas of densely packed nuclei and fewer human cells that colocalize with the dark spots in the T2w MRI (arrows in **d**). The inset shows the morphology of viable grafted cells. **g** Quantification of the volume of implanted COs from 8 mice. *b* indicates the animal shown in **b**. **h** Quantification of T2w intensity, normalized to the contralateral side, from 5 mice with trackable COs, showing a significant drop in T2w signals from D1 to D14 (*p* = 0.0002, paired t-test, n = 5 mice). **i** Diffusion Tensor Imaging (DTI) analysis of the CO engraftment site on D14. Top: T2w image with fluorescence overlay. Left: Apparent Diffusion Coefficient (ADC) map, showing spatial heterogeneity within the graft (white arrows). Right: Fractional Anisotropy (FA) map, revealing distinct signal enhancement at the graft-host interface (white arrowheads), suggesting axonal reorganization or ingrowth.

From the 8 mice that received CO transplantation, we drew regions of interests (ROIs) over the graft in T2w images. Relative to the mouse undergoing the same ischemic core removal procedure but without CO transplantation (**Supplementary Fig. 2a**), the implanted COs exhibited high T2w signal and this was confirmed by photos from surgical microscope (**Supplementary Fig. 2b**). We then quantified the volume of implanted COs ranges from 0.54 to 1.78 mm^3^, with a mean of 1.0 mm^3^ (**Fig. 3g**). T2 MRI images from the most representative planes on Day 1 from all animals included in this study are shown in **Supplementary Fig. 2b-c**. We noticed the T2w signal from the implanted CO on D14 (Fig. 3d, dashed region) is lower than that from D1 (Fig. 3b, D1, slice#4, dashed region). Quantification from 5 mice with trackable COs reveals a significant drop of T2w signals (**Fig. 3h**, *p* = 0.0001, n = 5 mice). This may be explained by the reduction in water content of the implanted CO, which may be a sign of the engraftment process.

As no clear borders of the CO graft could be revealed under T2w images at D14 post-transplantation, we performed diffusion tensor imaging (DTI) to generate apparent diffusion coefficient (ADC) and fractional anisotropy (FA) maps (**Fig. 3i**). Co-registration of fluorescence-based histology with T2w and DTI images enabled precise localization of the grafts within the infarct cavity. The indistinct graft-host boundary on T2w images suggests that the transplanted COs may have begun to integrate with the surrounding brain tissue. The ADC map showed spatial heterogeneity within the graft, likely reflecting variable cell density and extracellular matrix composition. Interestingly, out of the 22 parameters from the DTI analysis result (**Supplementary Fig. 3**), FA maps revealed a distinct signal enhancement at the graft–host interface, which lies beyond the graft into the host brain. This elevated FA signal, typically associated with aligned axonal structures, suggests axonal reorganization or ingrowth at the interface. Together, these findings demonstrate that DTI offers complementary, noninvasive readouts of graft maturation and host response—providing structural evidence of early tissue integration and potential neural remodeling induced by CO transplantation. Taken together, these MRI results establish a critical foundation for assessing transplantation efficacy in terms of anatomical integration and volumetric preservation, complementing subsequent functional and cellular imaging analyses.

### Longitudinal tracking of the viability of transplanted COs through BLI

To enable longitudinal, noninvasive assessment of graft viability following CO transplantation, we incorporated BLI as a core component of our MIPOT system. HiPSCs were transduced with a lentiviral construct encoding luciferase (Luc2) and mCherry, followed by monoclonal selection and differentiation into COs for transplantation to mice 10 days after the onset of photothrombosis (**Fig. 4a**). Luminescence reporter system allows longitudinal tracking of CO viability through BLI without the need for invasive procedures. To validate reporter expression, we monitored COs at early (3-day) and mature (3-week) stages of development. Compared to control organoids, Luc2+ COs showed strong mCherry fluorescence (**Fig. 4b**) and luminescence (**Fig. 4c**) at both time points. Quantification of luminescence confirmed significantly elevated signal in Luc2+ COs versus controls (*p* = 0.0002, **Fig. 4c**), with comparable diameters across conditions and time points (**Fig. 4d**, *p* = 0.0769, 3 days and 0.0946, 3 weeks). This indicates that the genetic modification associated with lentiviral labeling did not affect growth or differentiation potential of iPSCs as immunostaining COs with Tuji1 at older stages confirms that Luc2+ COs are capable of further neuronal maturation (**Supplementary Fig. 4a**).

**Fig. 4:**
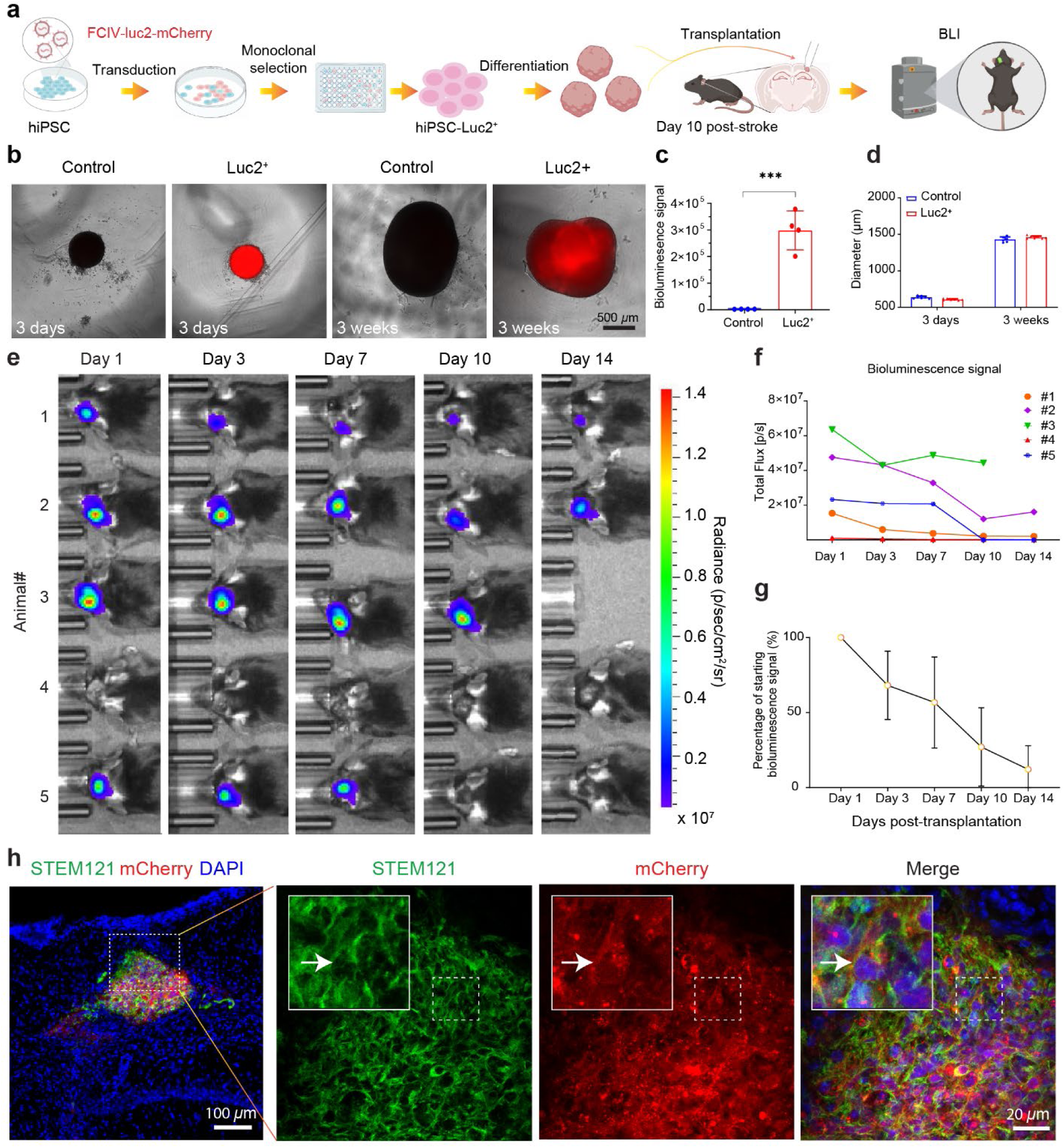
In vivo tracking of hiPSC-derived cortical organoid transplantation using bioluminescence imaging (BLI). **a** Schematic workflow of generating Luc2-mCherry dual-reporter hiPSCs, differentiation into COs, transplantation into stroked mouse brains at 10 days post-injury, and longitudinal tracking using BLI. **b** Brightfield and fluorescent images showing control and Luc2+ cortical organoids at day 3 and week 3 in vitro. Luc2+ organoids exhibit strong mCherry fluorescence without affecting gross morphology. **c** Quantification of luminescence signal confirms significantly higher signal in Luc2+ organoids compared to control organoids (\*\*\**p* = 0.0002, n = 4 COs per group, unpaired t test). **d** Diameter measurements of control and Luc2+ organoids at 3 days and 3 weeks show no significant difference, indicating stable growth kinetics. **e** In vivo BLI images from 5 animals over 14 days post-transplantation show localized signal from transplanted Luc2+ COs. **f** Longitudinal quantification of total BLI signal per animal shows a gradual decline over time in all animals except one, indicating partial but persistent viability. **g** Mean percentage of retained bioluminescent signal relative to day 1 reveals ∼25% of initial signal remains by day 14 (n = 4 animals). **h** Immunofluorescence analysis at endpoint (day 14) confirms survival of grafted cells. Human-specific STEM121 (green) and mCherry (red) co-expression within the graft (arrow) validates the in vivo BLI signal and confirms survival of transplanted human cells. Nuclei counterstained with DAPI (blue). All scale bars are as indicated in each panel.

Following transplantation into the ischemic cavity after wound cleaning at the chronic phase of stroke (10 days post-stroke), graft viability was monitored over two weeks using in vivo BLI (**Fig. 4e**). All animals (n = 5 mice) except 1 (mouse #4) exhibited localized above-background photon emission from the graft site beginning from day 1 post-CO transplantation. However, BLI signals declined progressively till 10 days post-transplantation and stabilized (**Fig. 4f**), with only ∼25% of the initial signal remaining by day 14 (**Fig. 4g**), indicating partial but sustained graft viability during early post-transplantation phase. We then performed histological analysis at the endpoint to confirm the presence of human cells within the host brain. Immunofluorescence for human-specific STEM121 (green) revealed grafted CO cells colocalized with mCherry (red) in the graft, validating the accuracy of the BLI signal and confirming survival of a subset of transplanted cells (**Fig. 4h** and **Supplementary Fig. 4b**). These results demonstrate that BLI within the MIPOT platform offers a sensitive, noninvasive approach to dynamically assess early graft viability after CO transplantation into the chronically ischemic brain.

### Labeling and high-resolution two-photon imaging iPSC-COs

To enable high spatial resolution tracking of grafted COs in vivo, we previously established a multicolor fluorescence labeling strategy and integrated it with two-photon fluorescence microscopy (TPFM), which is widely used for high-resolution in vivo brain imaging^28–30^. Our goal here is to monitor both the migration and functional activity of human iPSC-derived COs after transplantation into the ischemic cortex at the chronic phase of stroke. To achieve this, we utilized the RGB cell marking system, previously developed based on additive color mixing principles to distinguish individual cells by unique fluorescent hues (**Fig. 5a**), and recently optimized it with our newly developed ICam vector system for functional labeling of neurons ^27,31^. We constructed three ICam vectors, each co-expressing a distinct fluorophore—mCherry, TagBFP, or iRFP-682—linked via a self-cleaving 2A peptide to H2B-GCaMP6s (H2B-G66), a nuclear-localized calcium indicator (**Fig. 5b**). COs were labeled using either single color mode, where one cytosolic fluorophore was introduced (**Fig. 5b-c**), or mixed color mode, where a combination of vectors was used to generate a spectrum of color-coded cells in 7 patterns (**Fig. 5b**, right). In the single-color mode, wide-field fluorescence imaging showed distinct red and green fluorescent nuclei within the COs labeled with ICam-mCherry, confirming efficient gene delivery and expression (**Fig. 5c**). However, under wide-field fluorescence microscope, due to the scattering nature of the CO, it is challenging to obtain high-resolution images (**Supplementary Fig. 5a**). In contrast, TPFM enables subcellular resolution imaging of the live CO (**Fig. 5d**), revealing the nucleus-located G6s and cytosolic localization of mCherry (arrows in **Fig. 5d**). Strong second-generation harmonic (SHG) signal was detected in the live CO (**Fig. 5d**, arrow heads in the inset). In the mixed color mode, individual cells within the same organoid could be uniquely identified by their color hues from each of the three cytosolic FP and the nuclear G6s as revealed by the presence of colorful cells in the CO transduced with a 1:1:1 ratio mixture of the ICam vectors in terms of infectious unit (**Fig. 5e**). Importantly, these FPs could also be unmixed by special acquisition settings under TPFM^27^, enabling multicolor labeling of CO for potential functional cell tracking (**Supplementary Fig. 5b**).

**Fig. 5:**
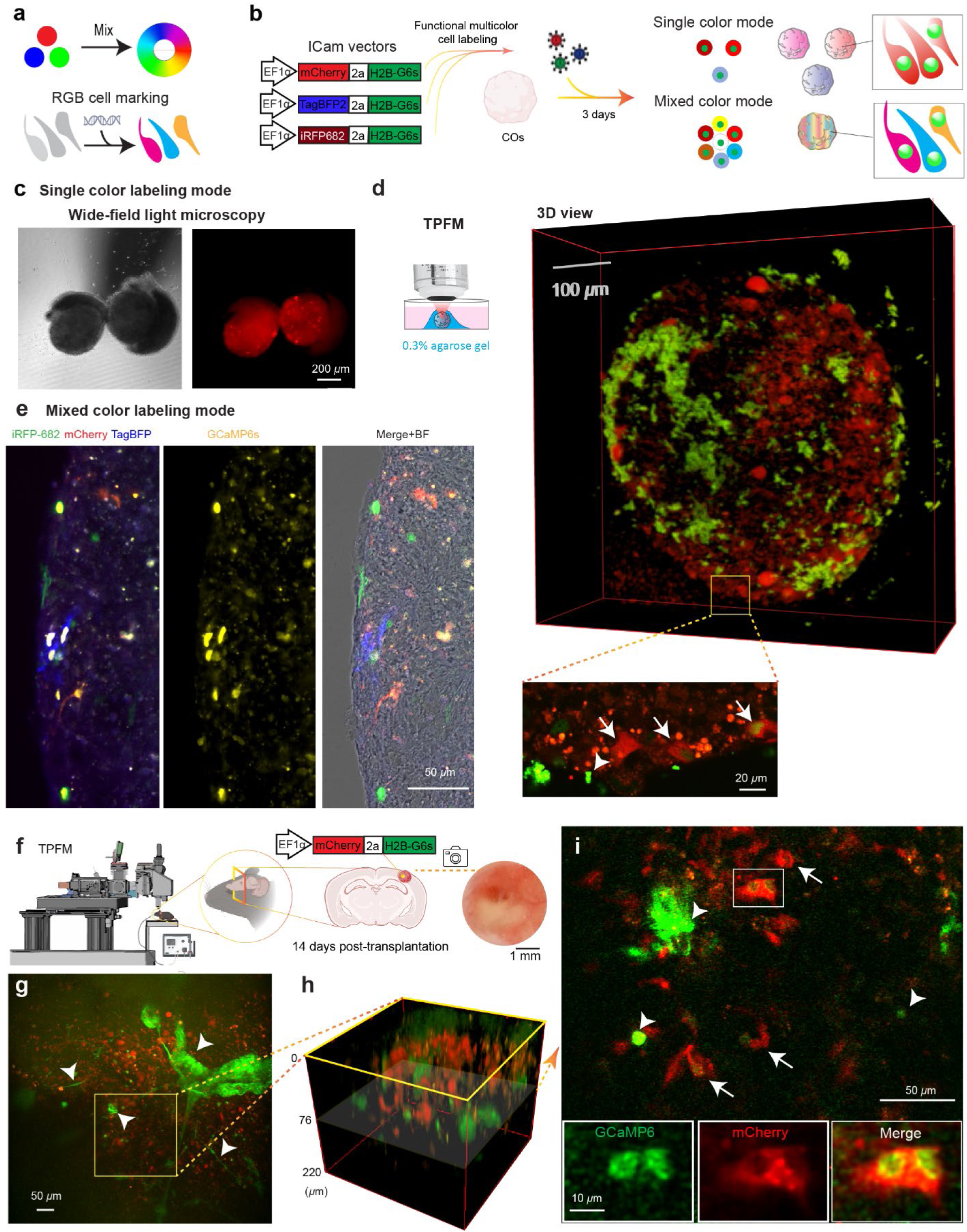
Fluorescent labeling of COs for TPFM. **a** Schematic of RGB cell labeling principle based on the combinatorial expression of three fluorescent proteins (mCherry, TagBFP, iRFP682), enabling a spectrum of colors via mixing. **b** Diagram of ICam vectors used for functional multicolor labeling of human iPSC-derived (COs). Three separate constructs drive the expression of nuclear H2B-GFP in tandem with mCherry, TagBFP, or iRFP682. Labeling results in single-color or mixed-color modes after 3 days of lentiviral transduction. **c** Wide-field bright field and fluorescence images of COs labeled in single-color mode by ICam-mCherry. **d** TPFM imaging of a CO labeled in single-color mode by ICam-mCherry. Left: schematic of in vitro imaging setup. Middle: 3D rendering of the organoid. Right: higher magnification image highlights distinct cells expressing mCherry and G6s (arrows). SHG signals were indicated by arrowheads. **e** Multichannel fluorescence images of a sliced CO labeled in mixed-color mode, showing mCherry, iRFP682, TagBFP, and GCaMP6s signals. Merge+Brightfield (BF) image confirms multicolor labeling within individual cells. **f** Schematic of in vivo two-photon imaging of COs grafted into the post-stroke cortex. At 14 days post-transplantation, labeled COs are imaged through a cranial window. **g** Representative in vivo two-photon image showing mCherry-positive grafted cells (red) and GCaMP6s expression (green). Arrowheads indicate grafted cells. **h** 3D reconstruction of the grafted CO by zooming in a region in (**g**). **i** The horizontal plane from (**h**) showing spatial distribution of mCherry and GCaMP6s signals as well as structured SHG signals (arrow heads). Insets show magnified views of the boxed region with individual channels and merged images, highlighting structural features of transplanted cells.

To explore the feasibility of in vivo imaging CO integration after transplantation into the chronically infarcted cortex, we performed intravital TPFM imaging 14 days post-transplantation (**Fig. 5f**). COs were pre-labeled with ICam-mCherry at 7 weeks post-differentiation and grafted at 8 weeks post-differentiation to chronic stroke in a manner similar to previous sessions. Under TPFM, CO cells were clearly identifiable within the stroke cavity *in vivo* (**Fig. 5g-i**). Interestingly, different from the in vitro CO, the SHG signal exhibited thick fiber-like 3-dimensional structures (**Fig. 5g-i, arrows**). High-resolution imaging revealed co-expression of mCherry and GCaMP6s, confirming the persistence of functionally active grafted cells (**Fig. 5i**, arrows and insets). Together, this system—integrating RGB-based multicolor labeling, genetically encoded calcium indicators, and intravital two-photon imaging—provides a robust platform to dynamically monitor cell fate and activity following CO transplantation in stroke. This multimodal approach lays the foundation for spatiotemporal analysis of organoid integration, behavior, and therapeutic efficacy in the living brain.

### Survival and differentiation of implanted COs for chronic stroke therapy

Using the MIPOT platform, we tracked graft dynamics through BLI during the early post-transplant phase, revealing an initial decline and gradual stabilization trend in viability during the first two weeks (**Fig. 4e-g**). This is consistent with the vulnerable window finding for transplanted cells or brain organoids, as it has been reported that the graft survival rate stabilizes after approximately 2-weeks post-engraftment^32–34^. To extend these observations, we analyzed brains collected at 2- and 4-weeks post-transplantation. Immunostaining for the human-specific antigen STEM121 confirmed the continued presence of COs at both time points, indicating sustained survival of grafted tissue in chronic stroke conditions (**Fig. 6a-b**). Interestingly, when COs were transplanted into the hippocampus, a robust differentiation phenotype was observed at 4 weeks post-transplantation (**Fig. 6c-d**). This is evidenced by the significantly longer process in 4 weeks group compared to 2 weeks or CO transplanted in the cerebral cortex from either 2 weeks or 4 weeks group (2 weeks cortex vs 2 weeks hippocampus, *p* = 0.527, 4 weeks cortex vs 4 weeks hippocampus, *p* < 0.0001, 2 weeks cortex vs 4 weeks cortex, *p* = 0.0004, 2 weeks hippocampus vs 4 weeks hippocampus, *p* < 0.0001, **Fig. 6e** and **Supplementary Fig. 6**).

**Fig. 6:**
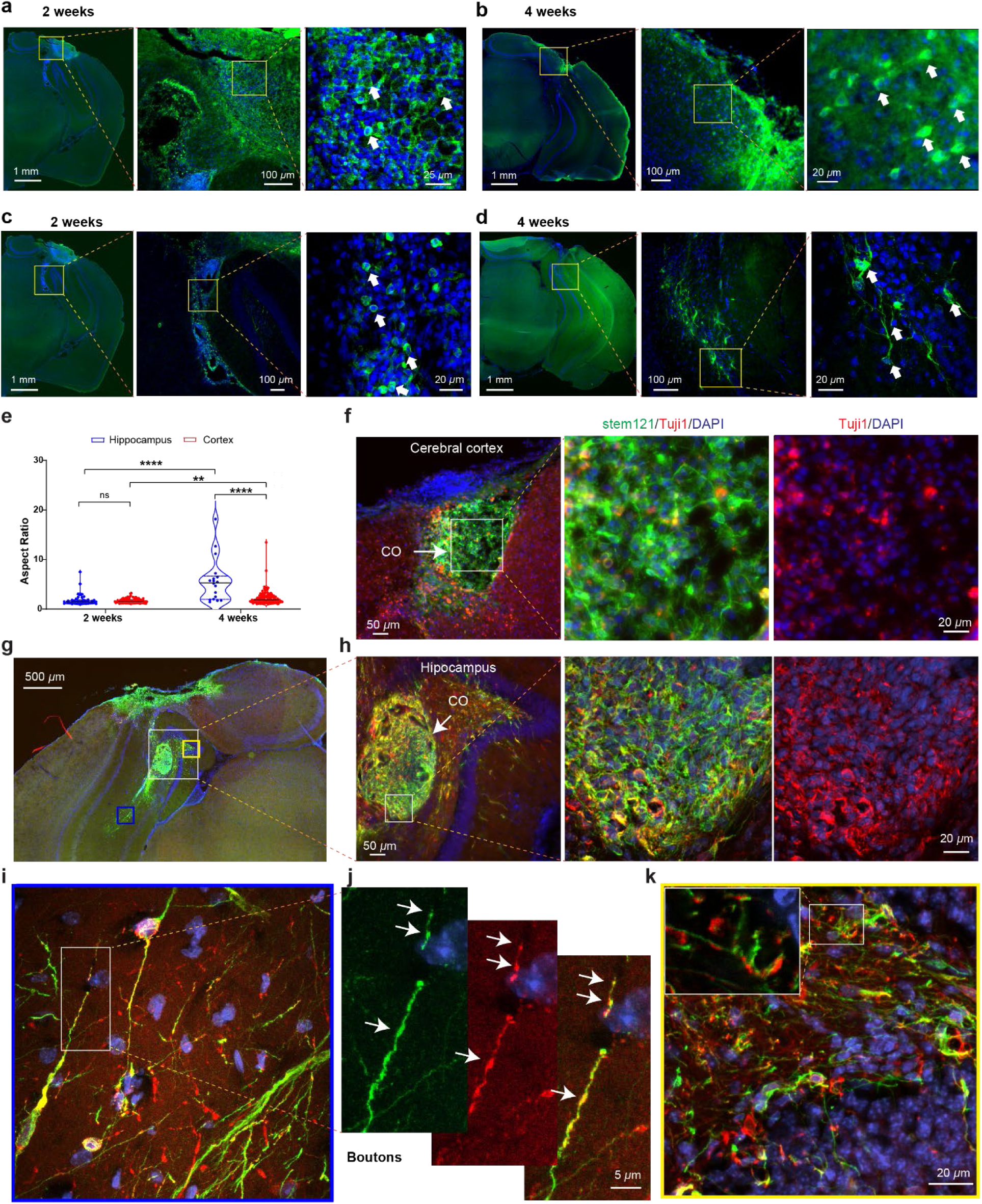
Region-dependent survival and neuronal differentiation of transplanted COs in chronic stroke mouse brains. **a–b**, Representative images of COs transplanted into the cerebral cortex at 2 weeks (**a**) and 4 weeks (**b**) post-transplantation. Immunostaining for human-specific marker STEM121 (green) demonstrates the persistent presence of grafted CO tissue. TUJ1 (βIII-tubulin, red) marks neuronal processes; nuclei are counterstained with DAPI (blue). **c–d**, COs transplanted into the hippocampus show greater morphological complexity at 4 weeks (**d**) compared to 2 weeks (**c**), including elongated processes (arrows), indicative of neuronal differentiation. **e**, Quantification of cell morphology in terms of aspect ratio to reflect process extension across transplant sites and time points. Cell counts were obtained from the cortex (2 weeks: *n* = 79 cells from 4 mice; 4 weeks: *n* = 121 cells from 5 mice) and hippocampus (2 weeks: *n* = 41 cells from 2 mice; 4 weeks: *n* = 18 cells from 2 mice). Cortex vs. hippocampus: 2 weeks (*p* = 0.527), 4 weeks (*p* < 0.0001); cortex 2 vs. 4 weeks (*p* = 0.0004); hippocampus 2 vs. 4 weeks (*p* < 0.0001). Statistical analysis was performed using the Mann–Whitney U test; \*\**p* < 0.01, \*\*\*\**p* < 0. 0001. **f**, In the cortex, a low percentage of TUJ1-positive cells (red) co-express STEM121 (green) at 4 weeks, suggesting delayed neuronal differentiation within cortical grafts. **g–h**, COs grafted into the hippocampus show robust co-expression of STEM121 and TUJ1 (yellow/orange) at 4 weeks. The graft core (CO) is well demarcated, with a high percentage of double-positive cells indicating enhanced neuronal commitment. **i**, At the graft periphery in the hippocampus, CO-derived cells extend long TUJ1-positive neurites with apparent integration into host tissue. **j**, High magnification of neurites showing bouton-like structures (arrows) co-labeled for STEM121 and TUJ1. **k**, In the dentate gyrus, processes of transplanted CO-derived cells (STEM121-positive) intercalate with host TUJ1-positive neurons, suggesting structural integration with the host network. All scale bars and magnifications are as indicated in each panel.

To assess neuronal differentiation of transplanted organoid-derived cells, we performed immunohistochemistry using the neuronal marker TUJ1 (βIII-tubulin). At 4 weeks post-transplantation, we observed low percentage of colocalization of TUJ1 with stem121 positive cells within the graft core in the cerebral cortex (12.8%, **Fig. 6f**), indicating neuronal differentiation of engrafted CO cells may take longer than 4 weeks, which is consistent with the lack of long process quantified in **Fig. 6e**. In stark contrast, COs cells grafted into the hippocampus exhibited high colocalization percentage with STEM121 (80.8%, **Fig. 6g-h**). Importantly, in the periphery of grafted CO, STEM121 and TUJ1 double-positive cells exhibited dense neurite processes, including axonal terminals (**Fig. 6i-j**). In the dentate gyrus, the processes of STEM121-positive cells are interleaved with TUJI1 cells (**Fig. 6k**), suggesting integration with host tissue.

This finding suggests that the hippocampal microenvironment promotes maturation and lineage commitment more effectively than cortical sites. Furthermore, we noted that cells located within dense graft cores harbor shorter and less sophisticated processes (**Fig. 6h**), whereas those that migrated into host tissue exhibited more mature neuronal morphologies characterized by extended dendritic processes and axonal boutons (**Fig. 6i-j**). This highlights the importance of migration from organoid masses as a requirement for further maturation.

In sum, our results demonstrate that COs not only survive but also retain structural stability following transplantation into chronic stroke lesions. Moreover, site-dependent differentiation is evident: cortex engraftments maintain morphology, while hippocampal implants promote neuronal maturation—especially among cells that exit the graft mass.

## Discussion

In this study, we established a Multimodal Imaging Platform for Organoid Tracking (MIPOT) that enables longitudinal, multi-scale assessment of hiPSC-derived COs transplanted into chronic stroke lesions. By combining intraoperative microscopy, high-field MRI, BLI, and TPFM, MIPOT allows us to confirm precise graft placement and to monitor CO survival, structural integration, and interactions with the host over time. We observed that CO grafts filled the lesion cavity immediately after delivery but experienced a rapid viability decline over the first ∼10 days, with only around a quarter of the initial cell survival by week 2. This early decline is consistent with prior reports that vascularized organoids shrink during the first 10–14 days before host perfusion is established^32,35,36^. Importantly, by 4 weeks post-transplantation, we still detect human cells within the cortex, confirming engraftment. Remarkably, COs that had migrated into the host hippocampus exhibited substantially enhanced maturation: these cells developed long neurites and expressed neuronal markers robustly, far more than COs confined to the cortical infarct. This site-dependent differentiation underscores that the host microenvironment strongly biases graft fate^37^. Together, our work not only developed a new approach for peri-transplantation period tracking of organoids but also revealed new biology – namely, the differential maturation of organoid cells in distinct brain niches.

Chronic phase stroke is featured by the formation of the ischemic core surrounded by glial scar, which prevents the integration of implanted cells or tissues. Previous work has shown that photothrombotic lesions stabilize between 7–10 days, with glial scar formation, hemorrhage resolution, and lesion volume plateauing during this subacute-to-chronic transition^25,38,39^. We reason that by removing the well-formed ischemic core, we could provide the grafted CO a clean parenchyma surface for them to integrate. Using our previously designed transplantation approach, pulse-elevation^27^, we managed to obtain high success rate of CO transplantation (8 out of 8, **Fig. 2g**), confirmed by MRI. This is solid step forward as consistency and reliability has been a major challenge for the transplantation of brain organoids due to the finicky nature and the lack of well-documented method in many previously published papers. The efficacy of this ischemic-core-removal and ensuing CO transplantation approach is demonstrated by the imaging and histology results showing viable cells at 2 or 4 weeks post-transplantation (**Fig. 6a-b**). To our knowledge, we are the first to advocate the removal of ischemic core in chronic stroke and using the resulting cavity for the accommodation of organoids for stroke treatment. Previous work on transplanting CO for stroke treatment focused on the acute or subacute phase of stroke^24^ or directly inject the CO into the junction of the infarct core and the peri-infarct zone of stroke^23^.

Most previous approaches for observation of transplanted brain organoids rely on single modalities or terminal histology as readout of brain organoids transplantation to lesioned ^20–24^ or developing animal brains ^32,40^. In one elegant study, multimodal monitoring of transplanted COs in the mouse brain was performed by combining two modalities: TPFM and microelectrode arrays for longitudinal tracking of COs^41^. Our MIPOT approach combines four intravital imaging modalities: surgical microscopy, MRI, BLI, and TPFM, offering transformative advantages in scale, resolution, depth penetration, and repeatability. Importantly, this platform supports tracking of the graft across weeks, enabling the observation of delayed or progressive changes in graft development and host response.

High-field MRI provided a noninvasive window into graft anatomy. However, so far there have been no studies that have performed MRI monitoring of implanted brain organoids from the peri-transplantation period. In our hands, COs appeared as distinct T2w hyperintensities in the lesion cavity on Day 1 post-transplantation (**Fig. 3b**), unambiguously confirming successful delivery. Over 2 weeks, we observed a significant decrease in T2w signal (**Fig. 3h**), which likely reflects reduced water content and compaction as edema resolves. We also collected diffusion MRI (DTI) metrics at 2 weeks and found elevated fractional anisotropy at the graft–host interface (**Fig. 3i**), suggesting aligned fibers or regenerating axons^42,43^. These MRI readouts (anatomy, volume, diffusion) provide quantitative gauges of engraftment. Importantly, histology registered to MRI confirmed that T2w hypointensities corresponded to densely packed nuclei (**Fig. 3d–f**), which may be the result of inflammatory cell invasion of the graft^44^. In sum, MRI validated our surgical technique and gave early clues to tissue integration, demonstrating the power of combining anatomic imaging with histological correlation.

To noninvasively measure graft viability, we labeled COs with a luciferase reporter and performed serial BLI (**Fig. 4e**). This revealed that most photon emission originated from the transplant site, and that luminescence declined markedly within the first 10 days (**Fig. 4f–g**). By day 14 the signal had plateaued at around a quarter of its peak. This finding is consistent with a general “vulnerable period” for grafts: other studies have found that stem-cell survival often stabilizes after ∼2 weeks post-transplant^32,35,36^. Crucially, our histology confirmed that the remaining bioluminescent signal was indeed from viable human cells (STEM121+ and mCherry+; **Fig. 4h**). Thus, BLI in MIPOT provided a quantitative in vivo measure of graft survival over time. As a method, BLI is uniquely sensitive for tracking transplanted cells: it has been widely applied to follow cell therapy in live animals ^45–47^. Our data demonstrate that only a fraction of transplanted COs persist long-term in chronic stroke, emphasizing the need for strategies (e.g. neurotrophic or angiogenic support) to improve survival. In the context of our work, the BLI signal decline defines the early graft-stabilization window, which complements the anatomic changes seen by MRI and will inform timing of future interventions.

We further exploited MIPOT’s multiscale capability by implanting fluorescently labeled COs and performing intravital TPFM. Using an RGB viral labeling system co-expressed with a nuclear calcium indicator (called ICam^27^), we were able to tag individual CO cells with color and image them at high resolution through TPFM (**Fig. 5d-e**). TPFM at 14 days post-transplant revealed labeled graft CO cells in situ (**Fig. 5g–i**). Interestingly, we observed thick SHG signals in fiber structure within the graft, possibly originating from collagen. This could be an indication of extracellular matrix remodeling or vascularization, which has been previously reported in transplanted brain organoids to the mouse brain^32,36,41^. Importantly, we directly visualized graft cell morphology at subcellular resolution: nuclei exhibited the calcium indicator signal and cytosol the chosen fluorophore, confirming that transplanted neurons remained intact and potentially active (**Fig. 5i**). Thus, TPFM completes the imaging cascade: it allows us to link graft viability (from MRI/BLI) with cellular phenotypes and interactions in the living brain. These data also imply host remodeling around the graft; the prominent SHG signal suggests host-derived collagen or myelin deposition, a phenomenon worth investigating for its impact on graft integration.

The most striking finding beyond our imaging toolkit was the region-specific graft outcome. COs left within the cortical stroke core at 4 weeks remained largely immature, with few long processes and limited neuronal marker expression (TUJ1); by contrast, cells that had migrated into the hippocampus showed robust neuronal differentiation and extensive neurite outgrowth (**Fig. 6d, 6i** and **6k**). This clearly indicates that the host niche governs graft fate. It is well-established that neural grafts differentiate in patterns dictated by local cues^37^. In particular, the hippocampus is a highly permissive, neurogenic environment: adult hippocampal stem/progenitor cells naturally generate new neurons^48^. Consistent with this, we observed much faster maturation of organoid-derived cells in the hippocampal milieu. These findings echo reports that NSC-derived neurons attain mature electrophysiological properties more quickly in hippocampal slices than in vitro cultures^37,49,50^. Our data suggest that biochemical signals, extracellular matrix, and synaptic inputs unique to the hippocampus promote CO-derived neuronal differentiation, whereas the chronically injured cortex lacks these supportive factors. Understanding these microenvironmental influences could guide improvements in transplant design, for example, co-transplanting helper cells expressing growth factors as shown in our previous work^47^. Overall, our results demonstrate that chronic stroke lesions can indeed host surviving organoid tissue, but that engrafted cells thrive only where the local environment supports maturation.

Despite demonstrating the feasibility of multimodal imaging to track CO transplantation in chronic stroke, our study has several limitations. First, our transplantation method involved splitting organoids and delivering them through a thick glass pipettes similar to needle^40^. While this approach allows better cavity filling and potentially enhances graft-host interface integration— crucial for stabilizing the cavity and preventing bleeding—it may introduce greater physical stress to the cells, potentially reducing viability compared to transplanting intact organoids^35,41^. Future studies could directly compare these approaches to evaluate the trade-offs between anatomical fit and graft survival. Second, although we used cyclosporin A for immunosuppression based on precedent in recent literature^35^, long-term studies would benefit from using genetically immunodeficient animals^36,40,41^ to ensure consistent suppression of host immune responses over extended periods. Third, although this 4-week analysis window reveals crucial dynamics, longer-term studies (e.g., ≥12 weeks) are needed to evaluate synaptic connectivity, electrophysiological maturity, and behavioral impact. Fourth, future work should incorporate vascular imaging agents and functional reporters to assess graft perfusion and neuronal activity over time. Behavioral assays will be critical to link cellular outcomes with functional recovery. Finally, exploring strategies to increase graft differentiation in the cortex—such as microenvironment modulation or region-specific organoid patterning—may pave the way for broader clinical application.

In conclusion, MIPOT provides a comprehensive platform to noninvasively chart the fate of human brain organoids in vivo. By applying it to chronic stroke, we have shown how multimodal imaging can bridge scales and modalities to yield new insights: from MRI-verified graft placement to BLI quantitation of viability, to two-photon views of cellular integration. These capabilities set the stage for mechanistic studies of organoid repair strategies and underscore the potential of organoid transplantation as a therapy for chronic stroke.

## Materials and Methods

### Human induced pluripotent stem cells culture

Human induced pluripotent stem cells (hiPSCs) were cultured as previously described^51^, with minor modifications. Cells were maintained under feeder-free conditions in mTeSR™ Plus medium (STEMCELL Technologies, Canada, Cat#100-0276) on vitronectin-X (STEMCELL Technologies, Canada, Cat#7180–coated plates. hiPSCs were passaged every 4–6 days at a 1:10 ratio using 0.5 mM EDTA (ThermoFisher, 440 Cat#25200056). For reseeding, cells were plated in vitronectin-XF–coated 6-well plates, and 10 μM Y-27632 (ROCK inhibitor; STEMCELL Technologies, Canada, Cat#72304) was added to the mTeSR™ medium for the first 24 hours post-passage to enhance cell survival. All hiPSCs used in this study exhibited typical pluripotent morphology and marker expression, with no signs of spontaneous differentiation. All hiPSC-related procedures were approved by the university’s ethics committee and complied with established ethical standards for human stem cell research.

### Lentiviral transduction and clonal selection of hiPSCs

Human iPSCs were transduced with the lentiviral vector pLV-FCIV-luc2-cherry. Briefly, cells were dissociated using 0.5 mM EDTA and resuspended at a density of approximately 1 × 10⁵ cells in 0.5 mL mTeSR Plus medium supplemented with polybrene (5 μg/mL), ROCK inhibitor (Y-27632, 10 μM), and 10 μL of lentiviral particles. Cells were incubated overnight at 37 °C, after which the medium was replaced with fresh mTeSR Plus.Two days post-infection, cells were seeded by limiting dilution into three 96-well plates at a density of ∼150 cells per plate. Clones expressing the reporter (cherry fluorescence) were identified under a fluorescence microscope, and wells containing single fluorescent-positive colonies were selected. After 7 days of culture in 96-well plates, selected clones were expanded into 24-well plates for further propagation and validation.

### Generation of human Cerebral Organoids

Generation of human cerebral organoids (hCOs) was performed based on the previously reported protocols^52,53^ Briefly, feeder-free hiPSCs were dissociated using Accutase (Sigma, Cat#A6964) and seeded at 9,000 cells per well into low-attachment 96-well U-bottom plates in mTeSR plus supplemented with 20 μM Y-27632. From day 1 to 5, spheroids were cultured in Essential 6™ Medium (Gibco, Cat#A1516401) containing 5 μM dorsomorphin (Sigma, Cat#P5499) and 10 μM SB-431542 (Tocris, Cat#1614). On day 6, spheroids were transferred to neural medium (NM) consisting of Neurobasal-A (Gibco, Cat#10888) supplemented with B27 minus vitamin A (Invitrogen, Cat#12587010), GlutaMAX (Invitrogen, Cat#35050-061), and penicillin-streptomycin (Sigma, Cat#P4333), along with 20 ng/ml bFGF (Peprotech, Cat#100-18B) and 20 ng/ml EGF (Peprotech, Cat#100-15), and cultured on an orbital shaker at 60 rpm. On day 25, organoids were transferred to low-attachment 24-well plates and maintained in neural medium supplemented with 20 ng/ml BDNF (Peprotech, Cat#450-02) and 20 ng/ml NT-3 (Peprotech, Cat#450-03). Medium was refreshed daily or every other day as appropriate.

### Lentiviral Transduction of Human Cerebral Organoids

Lentiviral transduction of hCOs was performed as previously described^54^, with minor modifications. Briefly, day 40–45 hCOs were individually transferred into 1.5-ml Eppendorf tubes containing 100 µl of NM supplemented with 8–10 µl of high-titer lentiviral vector (LV-hygro-ef1a-iRFP682-nes-h2bG6s, LV-hygro-ef1a-Cherry-nes-h2bG6s, LV-hygro-ef1a-TagBFP2-nes-h2bG6s, ∼10⁹ TU ml⁻¹; Vector Builder). Organoids were incubated at 37 °C for 45 min in a humidified incubator. Following transduction, each organoid was transferred into a well of a low-attachment 96-well plate containing an additional 100 µl of fresh NM and incubated overnight at 37 °C, 5% CO₂. The next day, organoids were washed once with 1 ml of fresh NM and transferred into low-attachment 24-well plates containing 1 ml of fresh NM per well. Organoids were maintained in culture with medium changes every 2–3 days. Transgene expression was monitored by fluorescence microscopy beginning 3–5 days post-infection. Robust expression was typically observed between days 5 and 7 and continued to increase over the following week. Only organoids displaying widespread expression were selected for downstream applications, including transplantation within 14 days post-infection.

### Mice

All experimental protocols at the University of Maryland, Baltimore, were conducted according to the National Institutes of Health guidelines for animal research and approved by the Institutional Animal Care and Use Committee at the University of Maryland, Baltimore. Mice were group housed with littermates until craniotomy surgery, after which they were singly housed. Mice were maintained on a 12–12-h (6 a.m.–6 p.m.) light–dark cycle. 3-4 months old male C57BL/6J mice (25-30 g, Institutional VR breeding) were used in this study. All mice were anesthetized with isoflurane during surgery and live imaging assays in this study (Vetone, 4-5% for induction and 1.5% for maintenance).

### Photothrombotic stroke model

Focal cortical infarcts in the motor cortex were induced using the photothrombotic stroke model, as described by Labat-Gest and Tomasi^55^. Briefly, mice were anesthetized with isoflurane (4–5% for induction, 1.5% for maintenance) and secured in a stereotaxic frame (RWD, Sugar Land, TX). A midline scalp incision was made, and the connective tissue was carefully removed to expose the intact skull surface. The photosensitive dye Rose Bengal (100 mg/kg body weight) was administered via intraperitoneal injection. A fiber optic cable connected to a cold light source was positioned 2.5 mm lateral to the midline and 1.4 mm anterior to bregma, targeting the motor cortex based on stereotaxic coordinates. Five minutes after dye injection, the targeted area was illuminated with a 532 nm laser for 15 minutes to induce a localized stroke (∼1 mm diameter) in the left cortex. Following the procedure, mice were returned to their cages and provided with moist food to support post-operative feeding.

### Organoid transplantation

CO transplantation was performed based on previously described protocols^56,57^ with slight modifications. Briefly, 10 days after stroke induction, a circular craniotomy (3 mm in diameter) was performed by drilling into the skull over the infarcted area, and the lesion was aspirated using vacuum suction to expose the cavity. One cerebral organoid (∼2 mm in diameter) was transferred onto a sterile parafilm, and excess medium was carefully removed. The organoid was then quartered along the midline using a sterile surgical blade. A 250-μm diameter glass pipette with a 45° beveled tip, connected to a 10-μL Hamilton syringe, was used for transplantation. One quarter of the organoid was gently loaded into the distal tip of the pipette. The syringe was mounted onto a microsyringe pump coupled with a stereotaxic manipulator. To calibrate the injection depth, the pipette tip was gently aligned with the surface of the cavity opening so that it was flush with the cavity edge; this position was defined as z = 0 and used as the reference for all subsequent depth measurements. The organoid delivery method followed a previously established protocol with minor modifications^58^. The injection was performed at three discrete depths: 1 μL was delivered at a depth of 1 mm, followed by a 1-minute pause. The pipette was then retracted to 700 μm, where a second 1 μL was injected, again followed by a 1-minute pause. Finally, the remaining organoid volume was injected at 500 μm depth, followed by a 3-minute pause to allow for tissue accommodation. The pipette was then slowly withdrawn at a controlled speed of 0.2–0.5 mm/min to minimize tissue disruption. Throughout the surgical procedure, the lesion site was continuously irrigated with sterile saline to maintain a clear, blood-free cavity. Hemostasis was achieved using a sterile gelatin sponge, which also absorbed excess blood and fluid. Following organoid implantation, the graft was covered with either a 3.5-mm glass coverslip or a layer of sterile plastic wrap to establish a cranial window. The implant was sealed using surgical adhesive. The scalp was then either sutured closed or, alternatively, a custom-designed headbar was attached to the skull using dental cement to allow for future head fixation during imaging. Upon completion of surgery, animals received analgesia via intraperitoneal injection of carprofen (5mg/kg, RimadyI). Mice were placed in a temperature-controlled recovery chamber and subsequently returned to their home cages. Immunosuppression was administered daily following transplantation with cyclosporin A at a dose of 10 mg/kg^59,60^.

### Tissue Collection and Immunofluorescence Staining

Animals were deeply anaesthetized and transcranial perfused with phosphate-buffered saline (PBS; Gibco, Cat#18912), followed by 4% paraformaldehyde (PFA in PBS; Sigma-Aldrich, Cat#P6148). Brains containing transplanted organoids were post-fixed overnight at 4 °C in 4% PFA and cryoprotected in 30% sucrose (Sigma-Aldrich, Cat#S9378) in PBS for 48–72 h. Organoids were fixed separately in 4% PFA for 2 h at room temperature and cryoprotected in 30% sucrose for 24 h. Tissues were embedded in OCT compound (brains: Tissue-Tek OCT Compound 4585, Sakura Finetek; organoids: 1:1 mixture of 30% sucrose and OCT), and coronally sectioned using a cryostat (CryoStar NX50) at 30 μm thickness (brains) or 20 μm (organoids). Cryosections were washed in PBS and incubated for 1 h at room temperature in blocking solution containing 10% normal bovine serum albumin (BSA; Sigma-Aldrich, Cat#A9647) and 0.3% Triton X-100 (Sigma-Aldrich, Cat#SLCD3084) in PBS. Sections were then incubated overnight at 4 °C with primary antibodies diluted in blocking solution. The following primary antibodies were used: anti-Stem121 (mouse, 1:500; Y40410, Takara Bio), anti-SOX2 (rabbit, 1:300; AB 5603, Millipose), anti-Nanog (rabbit, 1:200; ab 80892, Abcam), anti-CTIP2 (rat, 1:200; MABE1045, Sigma-Aldrich), anti-SATB2 (mouse, 1:50; ab51502, Abcam), anti-Tuj1 (rabbit, 1:200; 5568, Cell Signaling Technology), anti-Tuj1 (mouse, 1:500; 4466, Cell Signaling Technology), anti-TRA-1-60 (mouse, 1:100; MA1-023, Thermo Fisher), and anti-OCT4 (rabbit, 1:100; 11263-1-AP, Thermo Fisher). After washing with PBS, sections were incubated for 1 h at room temperature with fluorophore-conjugated secondary antibodies: Alexa Fluor 488 goat anti-rat (1:1000; A11006, Invitrogen), Alexa Fluor 488 goat anti-mouse (1:1000; A21121, Invitrogen), and Alexa Fluor 594 goat anti-rabbit (1:1000; A11012, Invitrogen). Nuclei were counterstained with Hoechst 33342 (1:10,000; 62249, Thermo Fisher). Finally, sections were mounted with Aquamount (F4680, Sigma-Aldrich) under cover glasses (Fisher Scientific) and imaged using either a Leica DMi8 fluorescence microscope or a Nikon A1 confocal microscope. Images were processed and analyzed using Fiji (ImageJ). To assess morphological changes of transplanted organoids over time in the cortical and hippocampal regions, the cells from grifted organoid boundaries were manually traced and measured using the “Freehand Selections” tool in Fiji.

### MRI acquisition and image analysis

Mice were scanned using a Bruker 9.4 T MRI system. T2-weighted images were acquired with a spin-echo sequence (TR/TE = 2500/33 ms) to assess the transplanted organoids at three time points: day −3 (7 days post-stroke), and day 1 and 14 post-implantation. Diffusion-weighted imaging (DWI) was performed across the brain using a spin-echo echo-planar imaging (EPI) sequence (TR/TE = 2300/25 ms, *b* = 650 s/mm²) with 30 diffusion encoding directions, following a previously published protocol^61^. A total of 11 slices (1 mm thickness each slice) were acquired at 14 days post-transplantation. Apparent diffusion coefficient (ADC) maps were generated to identify the stroke-affected slice and delineate stroke regions for subsequent region-of-interest (ROI) analysis. All other MRI scans were performed using single-slice acquisitions. Images analysis was conducted in Fiji (ImageJ). Organoid regions were manually delineated using the “Freehand Selections” tool and stored in the ROI Manager. Organoid volume was quantified only at day 1 post-transplantation using the formula: Volume = area × slice thickness, where area is the sum of segmented ROIs across relevant slices. T2 signal intensity was measured at both days 1 and 14 post-implantation. For normalization, anatomically matched contralateral regions were manually selected as reference ROIs. The signal intensity ratio between ipsilateral (organoid-containing) and contralateral regions was calculated to assess temporal changes in graft signal while controlling for inter-animal variability. DTI-derived images—including fractional anisotropy (FA), trace, signal intensity, trace-weighted images, tensor components (Dxx, Dyy, Dzz, Dxy, Dxz, Dyz), eigenvalues (λ1, λ2, λ3), and eigenvectors (X, Y, Z components of the first, second, and third eigenvectors)—were computed directly on the scanner. All datasets were exported using custom MATLAB scripts (MathWorks, Natick, MA, USA) for further quantitative analysis.

### BLI of CO grafts

Graft viability was longitudinally monitored using BLI. Mice were anesthetized with 2% isoflurane and intraperitoneally injected with D-luciferin (eLUCK-1g, Gold biotechnology) at a dose of 150 mg/kg body weight. Images were acquired every 5 minutes post-injection until the luminescence signal reached its peak intensity. For quantitative analysis, regions of interest (ROIs) were manually defined over the graft site, and total photon flux (photons/second, P/s) was calculated using Living Image software.

### Two-photon imaging and analysis

Mice were kept on a warm blanket (37 °C) and anesthetized using 0.5% isoflurane and Imaging was performed with a custom-built two-photon microscope with a resonant scanner. Each experimental session lasted 45 min to 2 h. Multiple sections (imaging planes) may be imaged within the same mouse. Fluorophores were excited by a different wavelength with a femtosecond laser system (Chameleon Discovery, Coherent) that was focused by an Olympus ×25, 1.05-NA objective. Emitted fluorescence photons reflected off a dichroic longpass beam splitter (FF705-Di01–25×36, Semrock) and were split with a dichroic mirror (565DCXR, Chroma) and detected by PMTs H16201P-40//004, Hamamatsu) after filtering with a 510/84-nm filter (84-097, Edmund) for the green channel and two 750SP filters (64-332, Edmund) for the red channel. Images were acquired using ScanImage^62^ (Vidrio Technologies). Three-dimensional z-stack images were acquired at a step size of 2 µm along the z-axis. After imaging, animals were returned to their home cages for recovery. In mice transplanted with labeled organoid, 1020 nm wavelength was used for mCherry and GcaMP6s. The laser power ranges from 4 to 16.8 mW.

Imaging data were processed with custom programs written in Fiji^63^. Raw two-photon imaging data were imported into ImageJ for the generation of composite images. Three-dimensional image reconstructions were obtained using the “3D Viewer. To enhance image contrast and resolution prior to quantitative analysis, we applied 2D deconvolution using the Richardson-Lucy (RL) algorithm for the image shown in Fig. 5d. The point spread function (PSF) was empirically measured from 3D image stacks of sub-diffraction fluorescent beads and averaged to obtain a representative system PSF. Image stacks were separated into their respective fluorescence channels, and each 2D frame was deconvolved independently using the averaged PSF. Deconvolution was carried out for 20 iterations using the RL algorithm, which is well established in fluorescence microscopy for mitigating spatial blurring caused by diffraction and optical aberrations.

### Statistical analysis

Data analysis was performed using a combination of standard functions and custom scripts in MATLAB, Prism 10 (GraphPad Software Inc. US) or PAST^64^. Data were tested for normality using the Shapiro–Wilk test. Parametric tests were applied to data that conformed to a normal distribution, whereas non-parametric tests were used for data that did not. Normally distributed data are presented as bar graphs showing the mean ± STD; non-normally distributed data are shown as box plots indicating the median and interquartile range (IQR). Box plots display the median and the 25th–75th percentiles, with whiskers drawn in the Tukey style (±1.5× IQR; Fig. 6e). For comparisons involving two factors, two-way ANOVA followed by Tukey’s multiple comparisons test was performed when data met the assumptions of normality (Fig. 4d). For comparisons between two groups, unpaired or paired parametric tests were used for normally distributed data (Fig. 4c and Fig. 3h, respectively), and the Mann Whitney test was used for unpaired, non-normally distributed data (Fig. 6e). Statistical significance was defined as follows: *\*\*p* < 0.01 *\*\*\*p*< 0.001 and *\*\*\*\*p* < 0.0001. Medians, IQRs, means, and SEM. values are reported throughout the text as appropriate.

## Acknowledgements

Y.L. acknowledges funding support from the National Institutes of Health (R21AG077631; R03NS123733; R03NS128459; R21AG074978) and Maryland Stem Cell Research Fund (2024-MSCRFD-6363 and 2022-MSCRFL-5893). P.W. acknowledges funding support from Maryland Stem Cell Research Fund (2022-MSCRFD-5886) and NIH (R01DA056739). J. W. acknowledges funding support from Maryland Stem Cell Research Fund for the postdoc fellowship (2024-MSCRFF-6328). T.-M.F. acknowledges funding support from the National Institutes of Health (1R21EB035681). We acknowledge the Confocal Microscopy Core Facility, and the Center for Translational Research in Imaging (CTRIM), supported by the Center for Innovative Biomedical Resources (CIBR) at the University of Maryland School of Medicine, for access to imaging instrumentation and expertise. Illustrative elements in this manuscript were created using academic resources from BioRender (www.biorender.com).

## Author contributions

Y.L. conceived the concept and designed the research. W.J. performed iPSC culture, differentiation, characterization, stroke modeling, transplantation, in vivo imaging, histology and data analysis. W.J. and G.Q. performed MRI scan. H.T. contributed to data analysis, stroke modeling and histology. M. J and P.W. contributed to the design and imaging of animal experiments. K.W. and J.X. contributed to MRI analysis. M.J.L and T.F. contributed to two-photon imaging analysis. Y. L. and P.W. supervised research. Y.L. and W.J. wrote the manuscript.

## Materials & Correspondence

All data are available from the Lead Contact, Yajie Liang, upon request.

## References

1. Tsao, C. W. et al. Heart Disease and Stroke Statistics—2023 Update: A Report From the American Heart Association. Circulation 147, e93–e621 (2023).

2. Hacke, W. et al. Thrombolysis with alteplase 3 to 4.5 hours after acute ischemic stroke. N. Engl. J. Med. 359, 1317–1329 (2008).

3. Saver, J. L. et al. Time to Treatment With Endovascular Thrombectomy and Outcomes From Ischemic Stroke: A Meta-analysis. JAMA 316, 1279–1288 (2016).

4. Duncan, P. W., Goldstein, L. B., Matchar, D., Divine, G. W. & Feussner, J. Measurement of motor recovery after stroke. Outcome assessment and sample size requirements. Stroke 23, 1084–1089 (1992).

5. Wechsler, L. R., Bates, D., Stroemer, P., Andrews-Zwilling, Y. S. & Aizman, I. Cell Therapy for Chronic Stroke. Stroke 49, 1066–1074 (2018).

6. Richards, C. L., Malouin, F. & Nadeau, S. Stroke rehabilitation: clinical picture, assessment, and therapeutic challenge. Prog. Brain Res. 218, 253–280 (2015).

7. Roth, E. J. Embracing change. Top. Stroke Rehabil. 22, 1 (2015).

8. Cramer, S. C. Recovery After Stroke. Contin. Minneap. Minn 26, 415–434 (2020).

9. Raza, S. S., Azari, H., Morris, V. B. & Popa Wagner, A. Editorial: Advances and challenges in stroke therapy: A regenerative prospective. Front. Neurosci. 16, 1102119 (2022).

10. Kadoshima, T. et al. Self-organization of axial polarity, inside-out layer pattern, and species-specific progenitor dynamics in human ES cell–derived neocortex. Proc. Natl. Acad. Sci. 110, 20284–20289 (2013).

11. Lancaster, M. A. et al. Cerebral organoids model human brain development and microcephaly. Nature 501, 373–379 (2013).

12. Qian, X., Song, H. & Ming, G.-L. Brain organoids: advances, applications and challenges. Dev. Camb. Engl. 146, dev166074 (2019).

13. Arlotta, P. Organoids required! A new path to understanding human brain development and disease. Nat. Methods 15, 27–29 (2018).

14. Kim, H. et al. Pluripotent Stem Cell-Derived Cerebral Organoids Reveal Human Oligodendrogenesis with Dorsal and Ventral Origins. Stem Cell Rep. 12, 890–905 (2019).

15. Lancaster, M. A. & Knoblich, J. A. Generation of cerebral organoids from human pluripotent stem cells. Nat. Protoc. 9, 2329–2340 (2014).

16. Hong, S. J., Bock, M., Zhang, S., An, S. B. & Han, I. Therapeutic Transplantation of Human Central Nervous System Organoids for Neural Reconstruction. Int. J. Mol. Sci. 25, 8540 (2024).

17. Wang, P. & Miao, C.-Y. NAMPT as a Therapeutic Target against Stroke. Trends Pharmacol. Sci. 36, 891–905 (2015).

18. Wei, L., Wei, Z. Z., Jiang, M. Q., Mohamad, O. & Yu, S. P. Stem cell transplantation therapy for multifaceted therapeutic benefits after stroke. Prog. Neurobiol. 157, 49–78 (2017).

19. Harary, P. M. et al. Cell Replacement Therapy for Brain Repair: Recent Progress and Remaining Challenges for Treating Parkinson’s Disease and Cortical Injury. Brain Sci. 13, 1654 (2023).

20. Jgamadze, D. et al. Structural and functional integration of human forebrain organoids with the injured adult rat visual system. Cell Stem Cell 30, 137–152.e7 (2023).

21. Bao, Z. et al. Human Cerebral Organoid Implantation Alleviated the Neurological Deficits of Traumatic Brain Injury in Mice. Oxid. Med. Cell. Longev. 2021, 6338722 (2021).

22. Zheng, X. et al. Human iPSC-derived midbrain organoids functionally integrate into striatum circuits and restore motor function in a mouse model of Parkinson’s disease. Theranostics 13, 2673–2692 (2023).

23. Cao, S.-Y. et al. Cerebral organoids transplantation repairs infarcted cortex and restores impaired function after stroke. Npj Regen. Med. 8, 27 (2023).

24. Wang, S.-N. et al. Cerebral Organoids Repair Ischemic Stroke Brain Injury. Transl. Stroke Res. 11, 983–1000 (2020).

25. Uzdensky, A. B. Photothrombotic Stroke as a Model of Ischemic Stroke. Transl. Stroke Res. 9, 437–451 (2018).

26. Labat-gest, V. & Tomasi, S. Photothrombotic Ischemia: A Minimally Invasive and Reproducible Photochemical Cortical Lesion Model for Mouse Stroke Studies. J. Vis. Exp. JoVE 50370 (2013) doi:10.3791/50370.

27. Wang, J. et al. OPTRACE: Optical Imaging–Guided Transplantation and Tracking of Cells in the Mouse Brain. bioRxiv 2025.05.24.655973 (2025) doi:10.1101/2025.05.24.655973.

28. Zipfel, W. R., Williams, R. M. & Webb, W. W. Nonlinear magic: multiphoton microscopy in the biosciences. Nat. Biotechnol. 21, 1369–1377 (2003).

29. Luu, P., Fraser, S. E. & Schneider, F. More than double the fun with two-photon excitation microscopy. Commun. Biol. 7, 1–15 (2024).

30. Liang, Y. & Walczak, P. Long term intravital single cell tracking under multiphoton microscopy. J. Neurosci. Methods 349, 109042 (2021).

31. Liang, Y. et al. Long-term in vivo single-cell tracking reveals the switch of migration patterns in adult-born juxtaglomerular cells of the mouse olfactory bulb. Cell Res. 26, 805–821 (2016).

32. Daviaud, N., Friedel, R. H. & Zou, H. Vascularization and Engraftment of Transplanted Human Cerebral Organoids in Mouse Cortex. eNeuro 5, (2018).

33. Reekmans, K. et al. Spatiotemporal evolution of early innate immune responses triggered by neural stem cell grafting. Stem Cell Res. Ther. 3, 56 (2012).

34. Tennstaedt, A., Mastropietro, A., Nelles, M., Beyrau, A. & Hoehn, M. In Vivo Fate Imaging of Intracerebral Stem Cell Grafts in Mouse Brain. PLOS ONE 10, e0144262 (2015).

35. Jgamadze, D. et al. Structural and functional integration of human forebrain organoids with the injured adult rat visual system. Cell Stem Cell 30, 137–152.e7 (2023).

36. Mansour, A. A. et al. An in vivo model of functional and vascularized human brain organoids. Nat. Biotechnol. 36, 432–441 (2018).

37. Kopach, O. Monitoring maturation of neural stem cell grafts within a host microenvironment. World J. Stem Cells 11, 982–989 (2019).

38. Li, H. et al. Histological, cellular and behavioral assessments of stroke outcomes after photothrombosis-induced ischemia in adult mice. BMC Neurosci. 15, 58 (2014).

39. Sunil, S. et al. Neurovascular coupling is preserved in chronic stroke recovery after targeted photothrombosis. NeuroImage Clin. 38, 103377 (2023).

40. Revah, O. et al. Maturation and circuit integration of transplanted human cortical organoids. Nature 610, 319–326 (2022).

41. Wilson, M. N. et al. Multimodal monitoring of human cortical organoids implanted in mice reveal functional connection with visual cortex. Nat. Commun. 13, 7945 (2022).

42. Jiang, Q. et al. MRI evaluation of axonal reorganization after bone marrow stromal cell treatment of traumatic brain injury. NMR Biomed. 24, 1119–1128 (2011).

43. Yamada, N. et al. Diffusion Tensor Imaging Evaluation of Neural Network Development in Patients Undergoing Therapeutic Repetitive Transcranial Magnetic Stimulation following Stroke. Neural Plast. 2018, 3901016 (2018).

44. Ghuman, H. et al. ECM hydrogel for the treatment of stroke: Characterization of the host cell infiltrate. Biomaterials 91, 166–181 (2016).

45. Huang, N. F., Okogbaa, J., Babakhanyan, A. & Cooke, J. P. Bioluminescence Imaging of Stem Cell-Based Therapeutics for Vascular Regeneration. Theranostics 2, 346–354 (2012).

46. Liang, Y., Walczak, P. & Bulte, J. W. M. The survival of engrafted neural stem cells within hyaluronic acid hydrogels. Biomaterials 34, 5521–5529 (2013).

47. Liang, Y., Ågren, L., Lyczek, A., Walczak, P. & Bulte, J. W. M. Neural progenitor cell survival in mouse brain can be improved by co-transplantation of helper cells expressing bFGF under doxycycline control. Exp. Neurol. 247, 73–79 (2013).

48. Bond, A. M., Ming, G. & Song, H. Adult Mammalian Neural Stem Cells and Neurogenesis: Five Decades Later. Cell Stem Cell 17, 385–395 (2015).

49. Morgan, P. J. et al. Human neural progenitor cells show functional neuronal differentiation and regional preference after engraftment onto hippocampal slice cultures. Stem Cells Dev. 21, 1501–1512 (2012).

50. Kopach, O. et al. Maturation of neural stem cells and integration into hippocampal circuits - a functional study in an in situ model of cerebral ischemia. J. Cell Sci. 131, jcs210989 (2018).

51. Li, W. et al. Rapid induction and long-term self-renewal of primitive neural precursors from human embryonic stem cells by small molecule inhibitors. Proc. Natl. Acad. Sci. U. S. A. 108, 8299–8304 (2011).

52. Sloan, S. A., Andersen, J., Pașca, A. M., Birey, F. & Pașca, S. P. Generation and assembly of human brain region–specific three-dimensional cultures. Nat. Protoc. 13, 2062–2085 (2018).

53. Paşca, A. M. et al. Functional cortical neurons and astrocytes from human pluripotent stem cells in 3D culture. Nat. Methods 12, 671–678 (2015).

54. Kelley, K. W. et al. Host circuit engagement of human cortical organoids transplanted in rodents. Nat. Protoc. 19, 3542–3567 (2024).

55. Labat-gest, V. & Tomasi, S. Photothrombotic ischemia: a minimally invasive and reproducible photochemical cortical lesion model for mouse stroke studies. J. Vis. Exp. JoVE 50370 (2013) doi:10.3791/50370.

56. Revah, O. et al. Maturation and circuit integration of transplanted human cortical organoids. Nature 610, 319–326 (2022).

57. Mansour, A. A. et al. An in vivo model of functional and vascularized human brain organoids. Nat. Biotechnol. 36, 432–441 (2018).

58. Wang, J. et al. OPTRACE: Optical Imaging-Guided Transplantation and Tracking of Cells in the Mouse Brain. BioRxiv Prepr. Serv. Biol. 2025.05.24.655973 (2025) doi:10.1101/2025.05.24.655973.

59. Kim, J.-T. et al. Human embryonic stem cell-derived cerebral organoids for treatment of mild traumatic brain injury in a mouse model. Biochem. Biophys. Res. Commun. 635, 169–178 (2022).

60. Wang, Z. et al. Cerebral organoids transplantation improves neurological motor function in rat brain injury. CNS Neurosci. Ther. 26, 682–697 (2020).

61. Wang, K. et al. Elucidating metabolite and pH variations in stroke through guanidino, amine and amide CEST MRI: A comparative multi-field study at 9.4T and 3T. NeuroImage 305, 120993 (2025).

62. Linaro, D. et al. Xenotransplanted Human Cortical Neurons Reveal Species-Specific Development and Functional Integration into Mouse Visual Circuits. Neuron 104, 972–986.e6 (2019).

63. Schindelin, J., et al. Fiji: an open-source platform for biological-image analysis. Nat. Methods 9, 676–682 (2012).

64. Hammer, O., Harper, D. A. T. & Ryan, P. D. PAST: Paleontological Statistics Software Package for Education and Data Analysis.

