## Supplementary Figures for "Intravital Multimodal Imaging of Human Cortical Organoids for Chronic Stroke Treatment in Mice"

**Supplementary Figure 1-6**

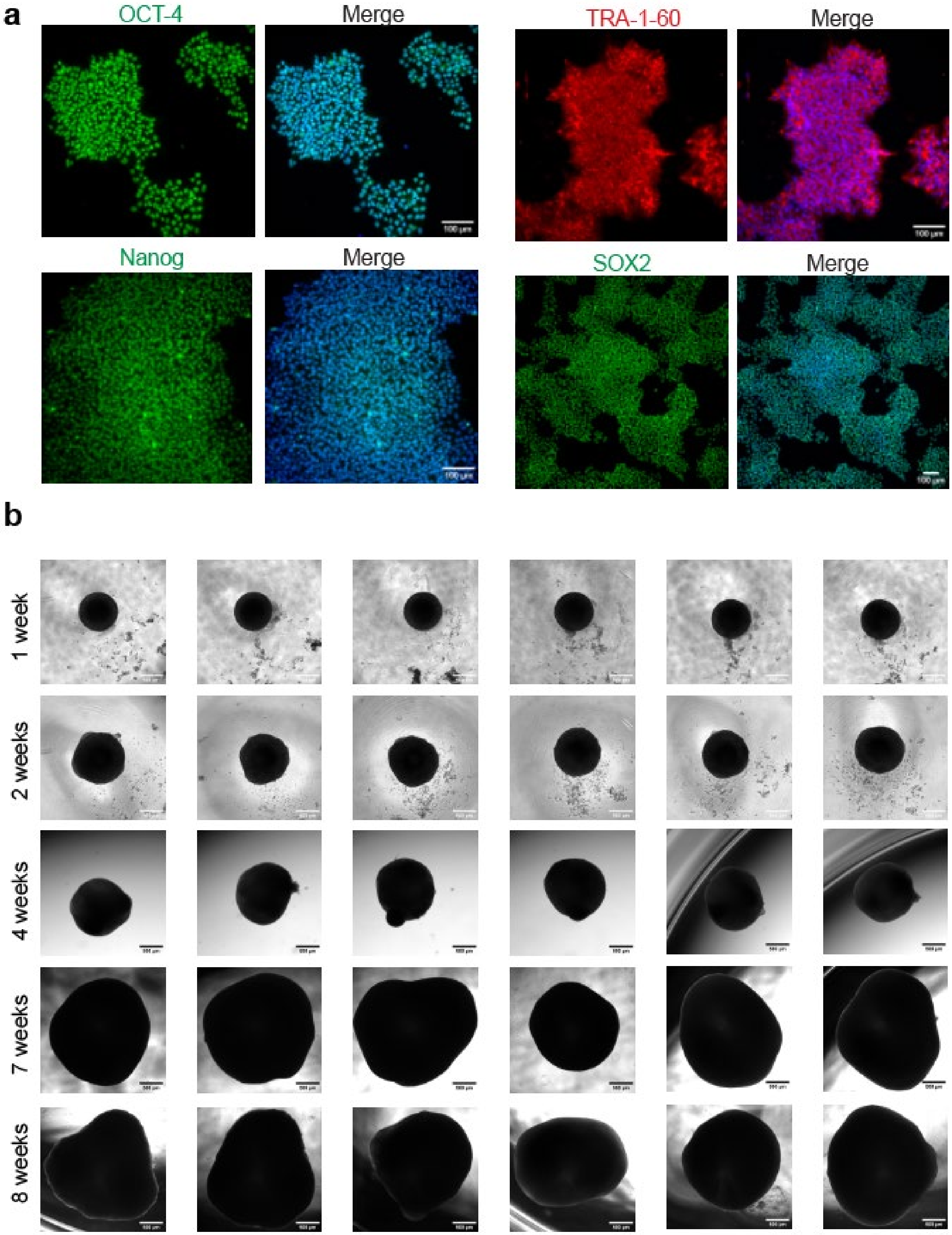
a
OCT-4
Merge
TRA-1-60
Merge
Nanog
Merge
SOX2
Merge
b
1 week
2 weeks
4 weeks
7 weeks
8 weeks

**Supplementary Figure 1. Characterization and long-term culture of hiPSC-derived COs.**

**a**, Immunofluorescence staining of pluripotency markers in human induced pluripotent stem cells (hiPSCs). Cells express OCT-4, TRA-1-60, Nanog, and SOX2, confirming maintenance of pluripotency. Nuclei were counterstained with DAPI. Scale bars, 100 μm.

**b**, Brightfield images of COs at 1, 2, 4, 7, and 8 weeks of differentiation. Organoids exhibit progressive growth and increased opacity over time, indicating tissue maturation. Each time point includes multiple representative organoids to show consistency across samples. Scale bars, 500 μm.

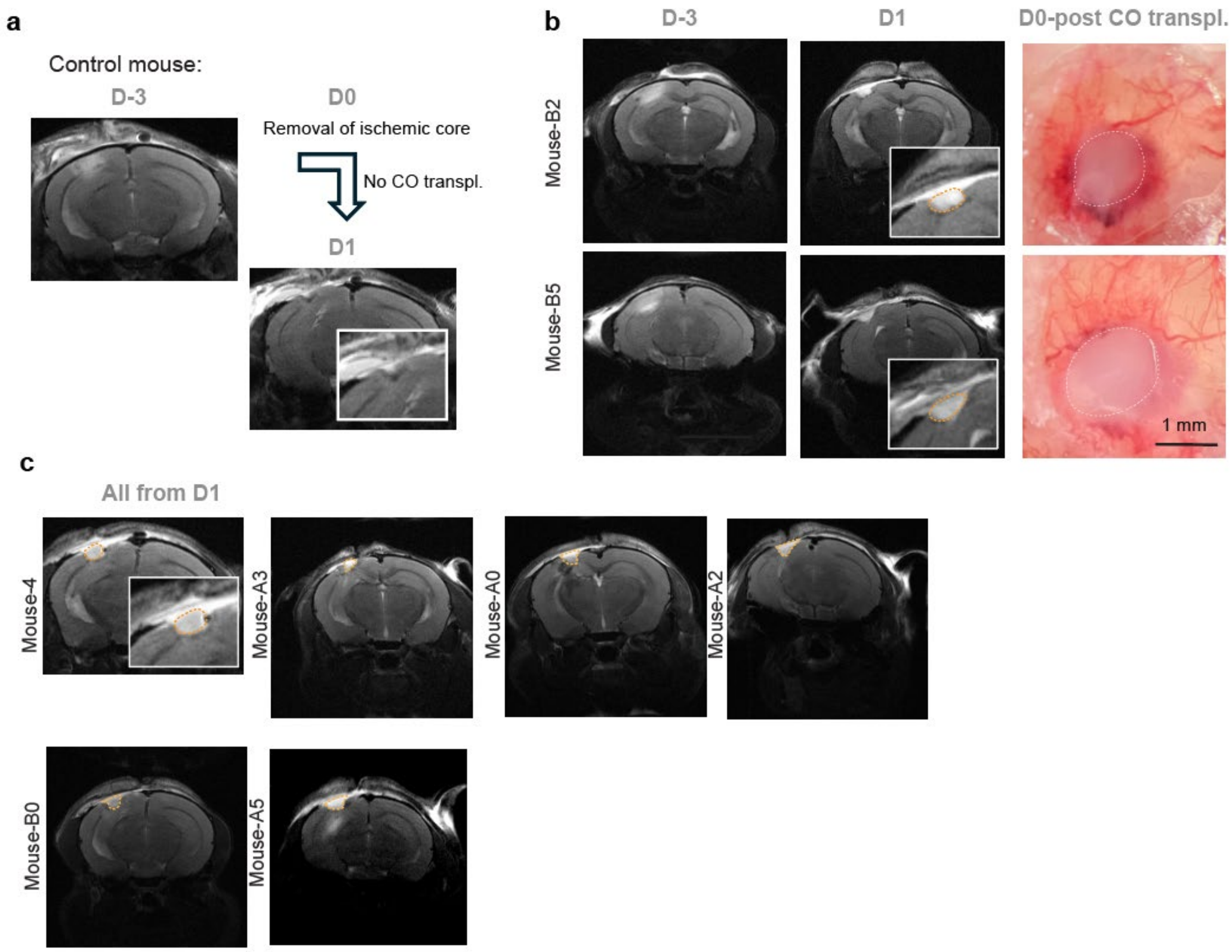


**Supplementary Figure 2. Confirmation of CO signal for quantification**.

**a**, T2-weighted MRI of a control mouse imaged by MRI at 3 days before surgery (D-3) and D1. The ischemic core was removed on D0, and no COs were transplanted. The inset at D1 shows the resulting cavity, which exhibits a low T2 signal.

**b**, Representative T2-weighted MRI images of two mice (Mouse-B2 and Mouse-B6) at D-3 and D1 post-transplantation of COs. The insets at D1 show the transplanted COs as distinct T2 hyperintensity regions within the infarct cavity. The right panel displays overhead surgical microscopy photos taken immediately after CO transplantation, confirming the presence of the graft (dashed white outline).

**c**, T2-weighted MRI images from the most representative planes on Day 1 from all animals (Mouse-4, Mouse-A3, Mouse-A0, Mouse-A2, Mouse-B0, Mouse-A5) included in the study for quantification in Fig. 3g, demonstrating the successful initial engraftment of COs. The insets show magnified views of the transplanted COs.

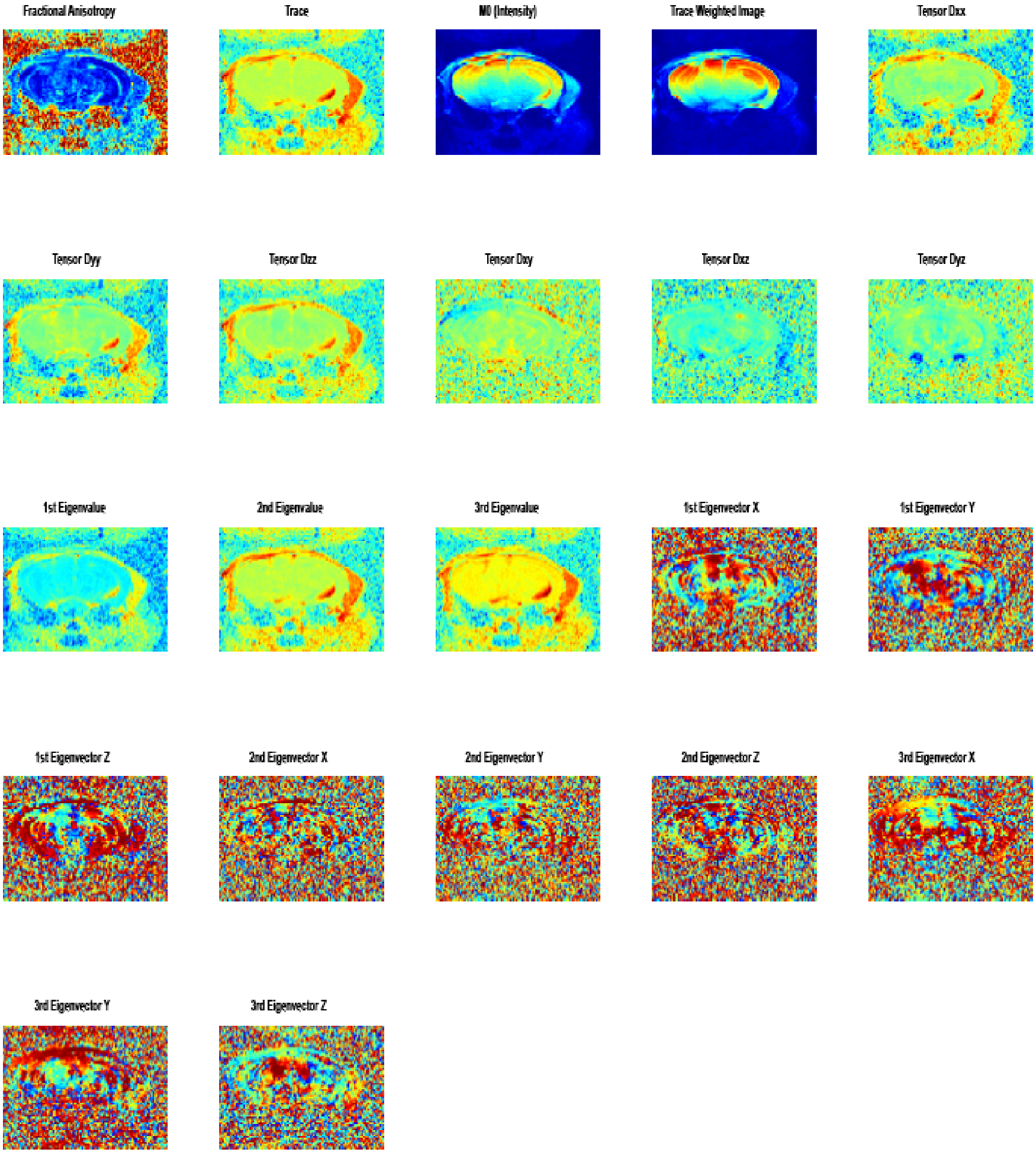


**Supplementary Figure 3. Diffusion tensor imaging analysis plots.** The 22 parameters plotted from the DTI analysis using jet as the color map.

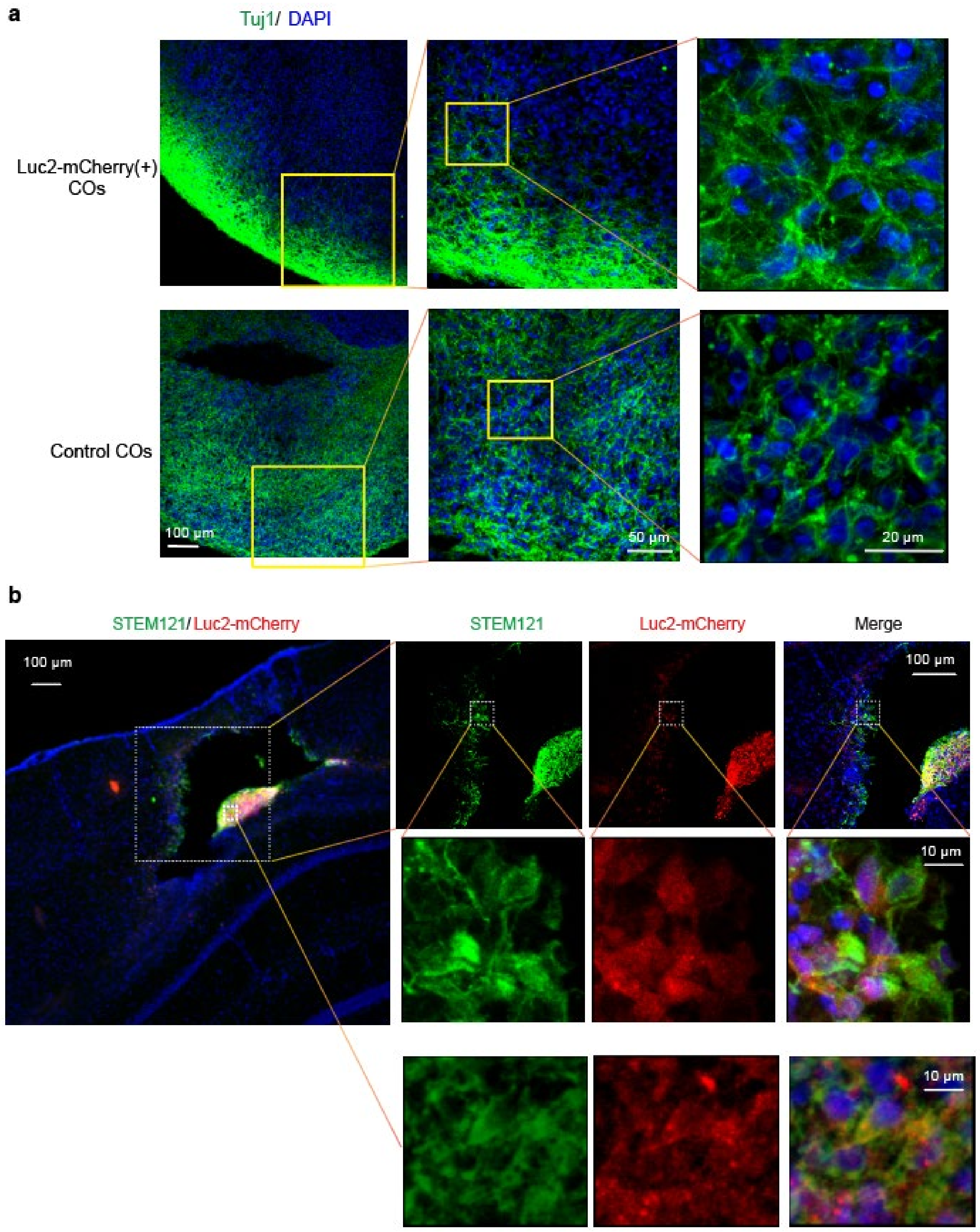


**Supplementary Figure 4. Validation of reporter expression and neuronal maturation in Luc2-mCherry–labeled COs.**

**a**, Immunofluorescence staining of COs derived from Luc2-mCherry–labeled hiPSCs (top) and unlabeled controls (bottom), showing neuronal marker Tuj1 (green) and nuclei (DAPI, blue). Representative images at increasing magnifications illustrate comparable neuronal differentiation between groups at the mature stage.

**b**, Immunostaining of grafted Luc2-mCherry(+) COs 14 days after transplantation into the stroke cavity. Human-specific STEM121 (green) colocalizes with Luc2-mCherry (red), confirming the survival and integration of donor-derived cells. Merged high-magnification images (rightmost panels) show cytoplasmic overlap of both markers.

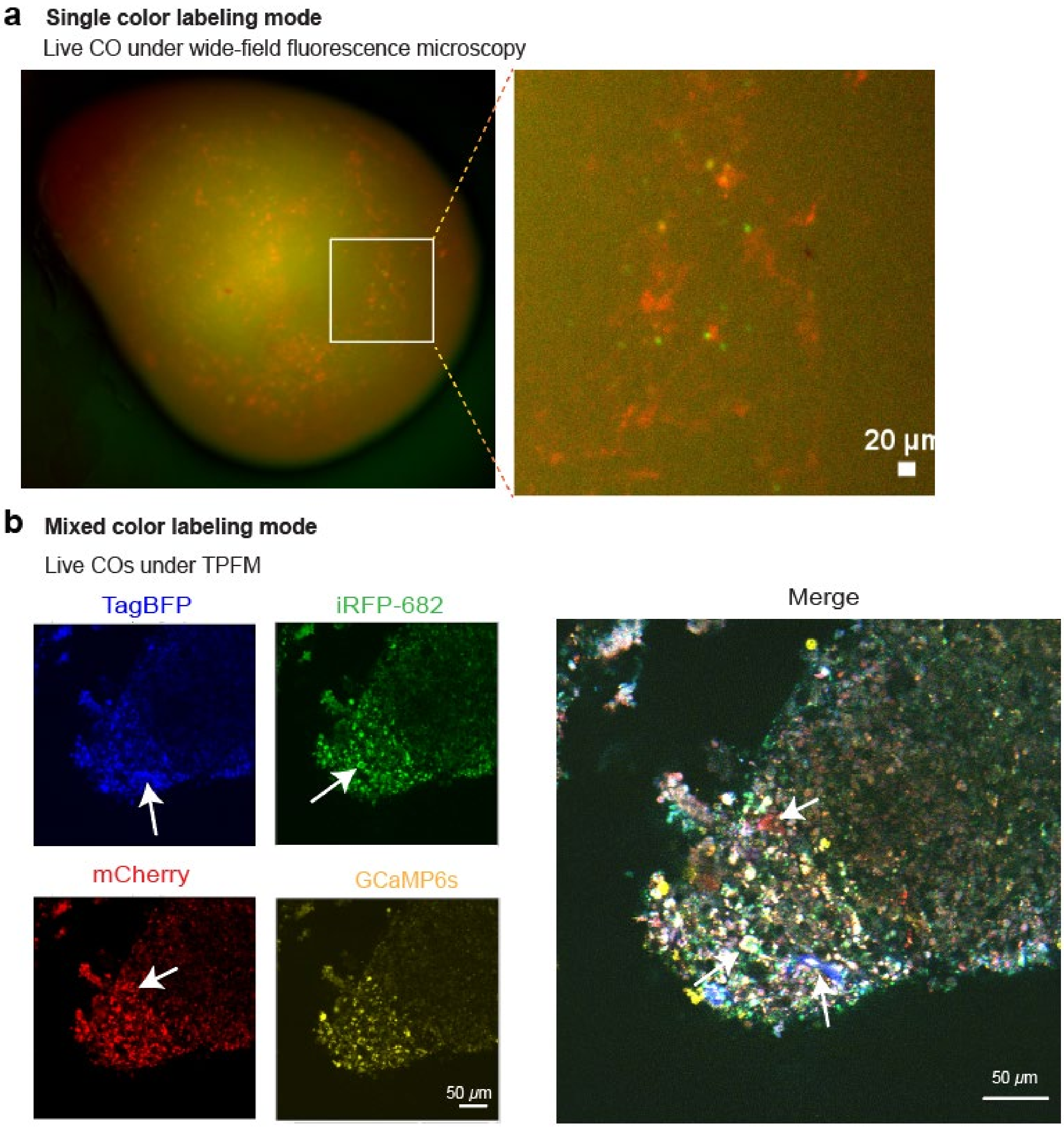


**Supplementary Figure 5. Multicolor labeling and high-resolution imaging of live COs using wide-field and two-photon fluorescence microscopy.**

**a,** Representative images of live COs labeled in single-color mode using ICam vectors expressing ICam:mCherry and GCaMP6s in the nucleus. Wide-field fluorescence microscopy revealed distinct mCherry (red) and iRFP-682 (green) expression within the CO, though spatial resolution is limited by tissue scattering.

**b**, Mixed-color labeling of live COs using a 1:1:1 ratio of ICam-mCherry, ICam-TagBFP, and ICam-iRFP-682 vectors co-expressing nuclear-localized H2B-GCaMP6s (G6s). TPFM allowed subcellular resolution of individual cells within the CO, each uniquely color-coded by fluorophore combinations (arrows). Multichannel acquisition enabled spectral separation of TagBFP (blue), iRFP-682 (green), mCherry (red), and GCaMP6s (yellow), with the merged image illustrating successful multicolor labeling. Each arrow points to a representative cell. This labeling and imaging approach provides a powerful tool to resolve spatial organization and monitor the behavior and function of individual CO-derived cells.

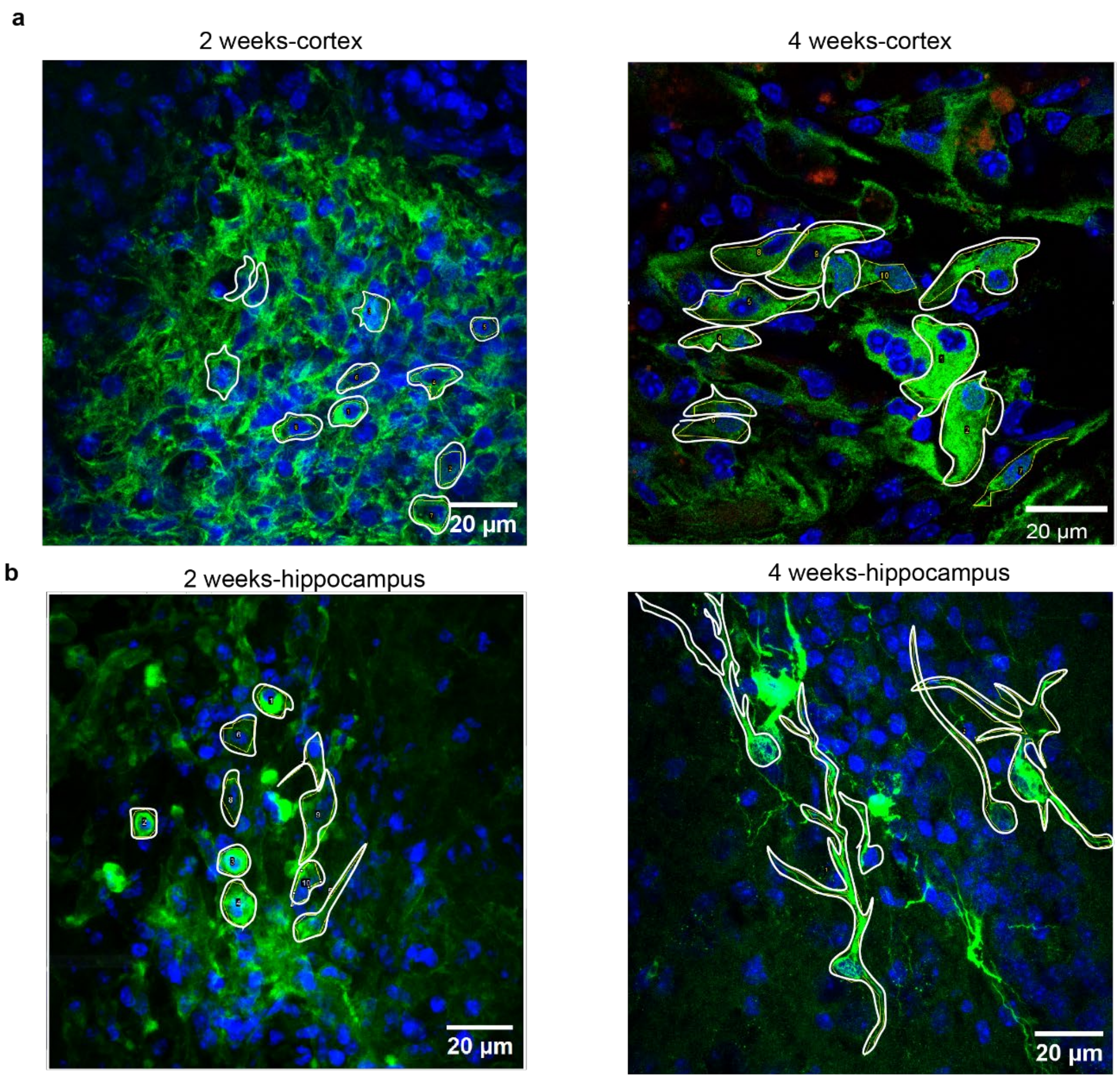


**Supplementary Figure 6. Examples of how ROIs were drawn on cells for quantification of aspect ratio.**

**a,** Brain slices containing CO in the cerebral cortex from 2-week or 4-week post-transplantation group. CO cells were outlined in white.

**b**, Brain slices containing CO in the hippocampus from 2-week or 4-week post-transplantation group. CO cells were outlined in white.
